# GLUD1 dictates muscle stem cell differentiation by controlling mitochondrial glutamate levels

**DOI:** 10.1101/2023.10.04.560525

**Authors:** Inés Soro-Arnáiz, Sarah Cherkaoui, Gillian Fitzgerald, Jing Zhang, Paola Gilardoni, Adhideb Ghosh, Ori Bar-Nur, Evi Masschelein, Pierre Maechler, Nicola Zamboni, Martin Poms, Alessio Cremonesi, Juan Carlos García Cañaveras, Katrien De Bock, Raphael J. Morscher

## Abstract

Muscle stem cells (MuSCs) enable muscle growth and regeneration after exercise or injury. Upon activation MuSCs metabolically rewire to meet the changing demands of proliferation. Here we describe that primary changes in metabolism itself can dictate MuSC fate decisions to control differentiation and fusion. We found that glutamine anaplerosis into the TCA cycle decreases during MuSC differentiation and coincides with decreased expression of the mitochondrial glutamate deaminase GLUD1. Genetic deletion of *Glud1* in proliferating MuSCs resulted in precocious differentiation and imbalanced fusion combined with loss of self-renewal *in vitro* and *in vivo*. Mechanistically, deleting *Glud1* caused mitochondrial glutamate accumulation in proliferating MuSCs and inhibited the malate-aspartate shuttle (MAS). Restoring MAS activity by supplementation of alanine normalized differentiation. In conclusion, high GLUD1 activity in proliferating MuSCs prevents deleterious mitochondrial glutamate accumulation and inactivation of the MAS. It thereby acts as a compartment specific metabolic brake on MuSC differentiation.

**Graphical Abstract:** 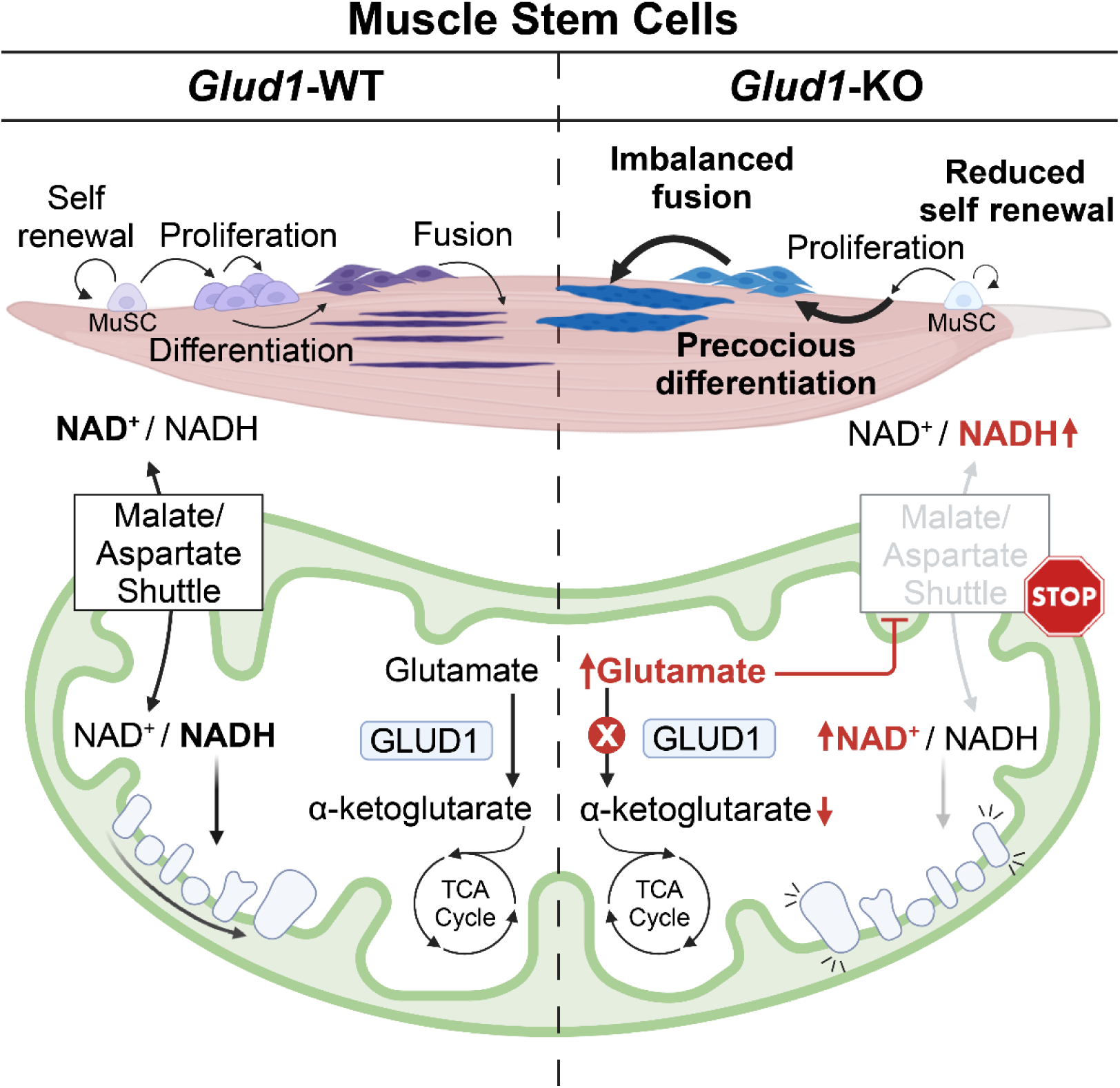

**Highlights:**

- Glutamine is the major TCA cycle substrate in MuSCs with decreasing contribution upon differentiation.
- Loss of *Glud1* impairs MuSC self-renewal capacity and causes imbalanced fusion *in vitro* and *in vivo*.
- *Glud1* deletion leads to mitochondrial glutamate trapping and malate-aspartate shuttle (MAS) dysfunction.
- Restoration of MAS activity in *Glud1* deficient MuSCs reverses precocious differentiation and imbalanced fusion.

## Introduction

Skeletal muscle possesses a remarkable capacity to adapt in response to training^1–3^ and regenerate upon injury^4,5^. Beyond the intrinsic plasticity of the muscle fiber, muscle has a unique microenvironment which supports adaptation and regeneration. Satellite cells, the muscle stem cells (MuSCs), which reside below the myofiber basement membrane, play a key role in muscle plasticity^6–8^. Under normal conditions, MuSCs are in a quiescent state and are characterized by the expression of the transcription factor PAX7^9–11^. Muscle injury or resistance training activates MuSCs to proliferate as myoblasts, and rapidly upregulate MYOD1^12–16^. Following several rounds of proliferation, a subset of MuSCs commits to differentiation, loses PAX7, and starts to express MYOGENIN^12,16^. Differentiated MuSCs contribute to muscle repair and growth by either fusing to pre-existing myofibers or to each other to form new myofibers. These fibers express mature myosin heavy-chain^17–20^. Importantly, an MuSC subset does not differentiate, but self-renews, and re-enters a quiescent state to maintain the PAX7^+^ MuSC pool^21–25^. The crucial role of MuSCs is underscored by genetic studies showing that these cells are necessary and sufficient for muscle regeneration^7,8^ and contribute to muscle hypertrophy^13,26,27^.

While under strong transcriptional control, MuSCs also undergo significant metabolic changes during myogenesis. Quiescent MuSCs have low metabolic activity and rely on glucose and fatty acid oxidation^28–31^. Upon activation, however, MuSCs rapidly rewire their metabolism to meet the energetic and biosynthetic demands of cell proliferation and growth. This includes increased glycolysis^29^ and mitochondrial biogenesis^32^, as well as the use of a broader substrate repertoire for aerobic energy production^29–31^. Consistent with a preferential breakdown of glucose to lactate, proliferating MuSCs have low pyruvate dehydrogenase (PDH) activity^31,33^. Nonetheless, glucose that enters the TCA cycle fuels cytosolic acetyl-CoA via export of citrate that increases histone acetylation to facilitates myogenic gene expression^31^. Histone acetylation is likely further enhanced by reduced SIRT1 deacetylase activity and lower NAD^+^ levels due to higher glycolysis^30^. Overall, muscle progenitors rewire their metabolism during activation and this reprogramming is essential for effective myogenesis and muscle regeneration.

Whereas metabolic changes associated with the transition from quiescent MuSCs to proliferative myoblasts have been actively studied, the metabolic rewiring associated with their differentiation to myotubes, is less understood^29–31,34–36^. First, data indicate that differentiating myoblasts and myotubes increasingly use PDH to shunt glucose carbons into the TCA cycle to fuel oxidative phosphorylation in the mitochondria. Also, the contribution of glucose to cytosolic acetyl-CoA, the substrate for nuclear acetylation reactions, is reduced^31,33^. The quantitative changes in metabolism affecting TCA cycle carbon sources during early and later steps of differentiation, however, are poorly described.

This is particularly true for amino acid metabolism. Through their role as regulators of mTORC1 signaling, amino acids crucially contribute to the exit from quiescence and prepare MuSC proliferation. Activation of mTORC1 in response to injury-induced systemic signals brings quiescent MuSCs into a G_alert_ state, an adaptive response to prime them for cell cycle entry^32^. Inhibition of glutamine uptake by deleting the glutamine transporter SLC1A5 reduces MuSCs proliferation *in vitro* and *in vivo* in a mTORC1 dependent manner^37^. At the same time, loss of *Slc1a5* suppressed muscle differentiation^37^, but the mechanism for the lack of glutamine import suppressing muscle differentiation is not clear. Overall, the metabolic fate of glutamine in MuSCs has not been thoroughly investigated.

After cellular uptake, glutamine can either be imported into the mitochondria^38^, its primary site of metabolism, or directly metabolized in the cytosol by asparagine synthetase (ASNS)^39–41^. In the mitochondrial matrix glutaminase liberates the amide-nitrogen by deamidation to form ammonia and glutamate. Depending on the cell’s enzymatic toolkit, glutamate is then either deaminated or serves in transamination reactions with alpha-ketoacids to form mitochondrial aspartate and alanine. Both reactions produce alpha-ketoglutarate (aKG), which contributes a five-carbon unit to the TCA cycle as an anaplerotic source^42^. Whether subcellular compartmentalization of glutamine metabolism affects MuSC differentiation is unclear.

Here, we used isotope tracing to characterize changes in metabolism of MuSCs following differentiation. We found that proliferating MuSCs use glutamine as their dominant TCA cycle substrate. Upon differentiation, glutamine anaplerosis to the TCA cycle reduces, coinciding with a lower expression of the mitochondrial deaminase GLUD1. Genetic deletion of *Glud1* in proliferating MuSCs resulted in precocious differentiation with loss of self-renewal *in vitro* and *in vivo,* combined with imbalanced fusion. Mechanistically, deleting *Glud1* caused mitochondrial glutamate accumulation in proliferating MuSCs and inhibited the malate-aspartate shuttle (MAS). Restoring MAS activity by supplementing MuSCs with alanine reverted imbalanced fusion. Our data show that GLUD1 activity in proliferating MuSCs ensures optimal glutamine anaplerosis, while preventing deleterious mitochondrial glutamate accumulation and inactivation of the MAS shuttle. We thereby highlight a crucial contribution of compartmentalized amino acid metabolism for MuSC fate decisions.

## Results

### Muscle stem cell differentiation is accompanied by a shift in anaplerotic TCA substrates

To elucidate how MuSCs rewire their metabolism during late myogenic differentiation, we isolated primary mouse MuSCs and compared their metabolism in a proliferative state (myoblasts) to a state of terminal differentiation (myotubes). Classic primary MuSC proliferation and differentiation media strongly differ in composition^27^, especially with respect to glucose and amino acid content (Table S1). To exclude the influence of media and isolate the cell intrinsic metabolic phenotypes, we established a differentiation protocol that only differs in the proportion of growth factors and dialyzed serum present in the media, while allowing normal differentiation (Figure 1A, S1A and S1B). Under those standardized culture conditions, we performed a global analysis of metabolite abundance in myoblasts and myotubes. Principal component analysis (PCA) showed clustering based on differentiation status with PC1 explaining more than 50 % of variation, suggesting distinct metabolic programs in myoblasts as compared to myotubes (Figure 1B). Pathway enrichment analysis of metabolites revealed TCA cycle as one of the top enriched pathways in myotubes (Figure 1C). Focusing on TCA cycle intermediates, we found that, while citrate abundance was increased, aKG levels were lower in myotubes when compared to myoblasts (Figure S1C). Moreover, the glutamate / aKG ratio increased in myotubes (Figure 1D), suggesting changed kinetics in the contributions from glutamine to the TCA cycle. Together, these pool size measurements indicated a reprogramming of the TCA cycle upon MuSC differentiation that might be related to changes in anaplerotic contributions from glutamine.

**Figure 1.**
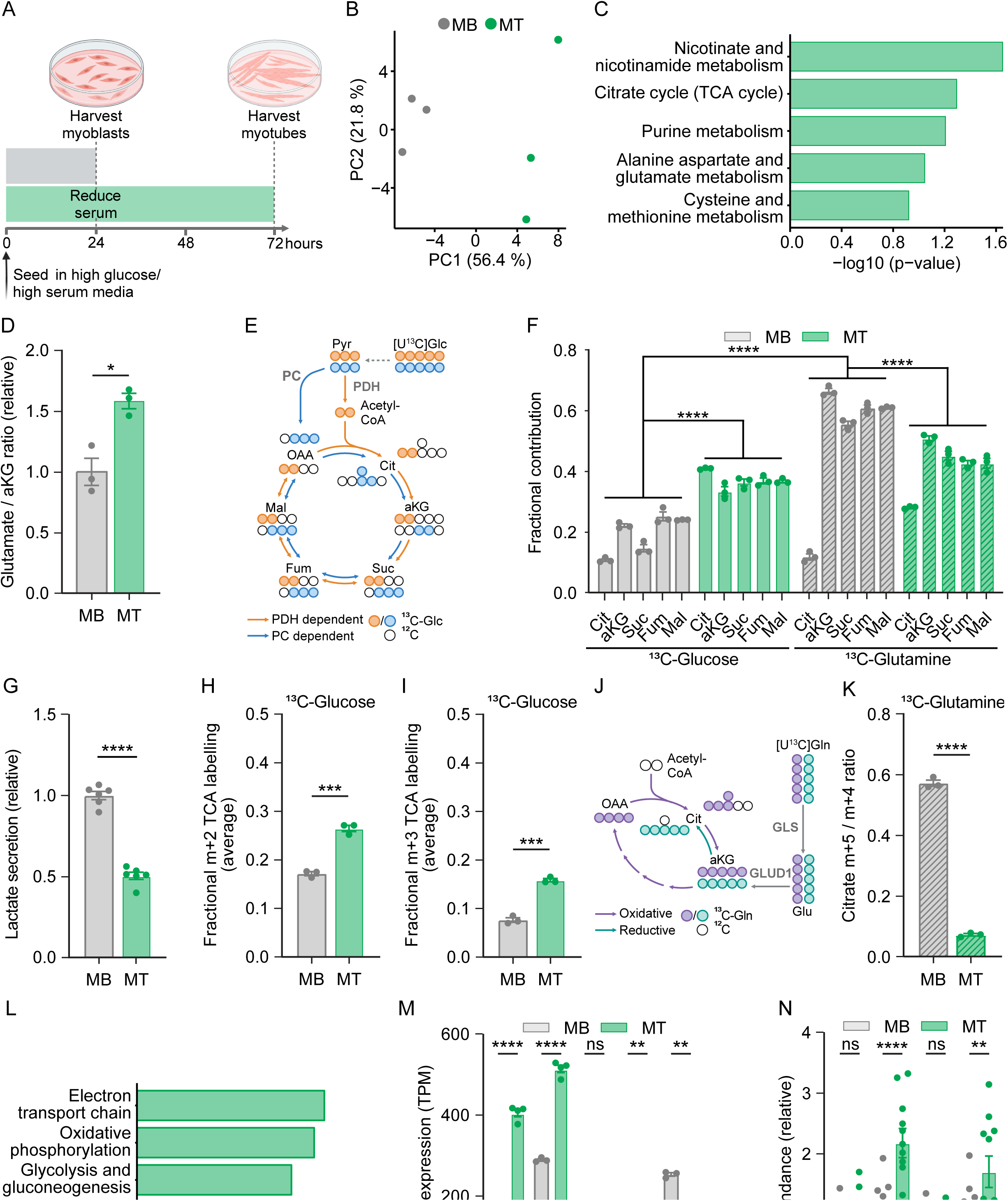
Muscle stem cell differentiation shifts substrate choice for TCA cycle anaplerosis. (A) Schematic representation of MuSC differentiation protocol where cells are seeded in high glucose / high dialyzed serum medium and differentiated in high glucose / low dialyzed serum medium. (B) Principal component analysis (PCA) from untargeted metabolomics of myoblasts (MB) and myotubes (MT). (C) Pathway enrichment analysis from untargeted metabolomics comparing MB to MT. Displayed are top 5 pathways from KEGG metabolic pathways showing differential regulation in metabolites levels. n = 3 per group. (D) Relative glutamate / alpha-ketoglutarate ratio in MB and MT (E) Schematic representation of tracing the contributions of [U^13^C]glucose to the TCA cycle through pyruvate carboxylase (PC, blue) and pyruvate dehydrogenase (PDH, orange). (F) Fractional contribution of [U-^13^C]glucose or [U-^13^C]glutamine to TCA cycle intermediates citrate (Cit), alpha-ketoglutarate (aKG), succinate (Suc), fumarate (Fum), and malate (Mal) in MB and MT. (G) Relative abundance of lactate in the media supernatant of MB and MT normalized to the packed cell volume. (H and I) Average fractional (H) m+2 labelling or (I) m+3 labelling from [U-^13^C]glucose to the TCA cycle intermediates citrate, alpha-ketoglutarate, succinate, fumarate, and malate in MB and MT (J) Schematic representation of [U^13^C]glutamine tracing in the TCA cycle through oxidative (purple) or reductive (turquoise) metabolism. (K) Citrate m+5 / m+4 labeling ratio in MB and MT cultured in media containing [U-^13^C]glutamine. (L) Top 5 upregulated pathways identified using gene set enrichment analysis with WikiPatways. Positive normalized enrichment score (NES) denotes higher expression in MT compared to MB. MB: n = 3, MT: n = 4. (M) Gene expression of the transaminases *Got1, Got2, Gpt1, Gpt2*, and the deaminase *Glud1* in MB and MT. (N) Relative abundance of the amino acids alanine (Ala), aspartate (Asp), glutamine (Gln), and glutamate (Glu) in MB and MT. Bar graphs represent mean ± SEM. Each dot represents a biological replicate. Student’s t test (two-tailed, unpaired) in (D), (G-I), and (K). Two-way ANOVA with Tukey’s multiple comparison test in (F). Two-way ANOVA with Sidak’s multiple comparison test in (M and N). (*p < 0.05, **p < 0.01, ***p < 0.001, ****p<.0001)

To explore the actual carbon flow to the TCA cycle we performed stable isotope tracing in myoblasts and myotubes. First, the metabolic fate of glucose was followed by culturing them in the presence of [U-^13^C]glucose (Figure 1E). This revealed an increased fractional contribution of glucose to all TCA cycle intermediates (Figure 1F, left), upon differentiation. Concomitantly, lactate secretion was reduced (Figure 1G), supporting a switch from anaerobic glycolysis to glucose oxidation^31^. The contribution to TCA cycle was characterized by a significantly higher m+2 labelling fraction in myotubes, consistent with enhanced PDH activity upon differentiation^31^ (Figure 1H and S1D). Besides this increased glucose contribution to the TCA cycle via acetyl-CoA, we also detected a higher m+3 labeling fraction of all intermediates (Figure 1I and S1D). This provides evidence that glucose-derived carbons are also increasingly used to replenish the TCA cycle via pyruvate carboxylase (PC). Given the large missing fraction in carbon source upon glucose tracing (Figure 1F, left), we investigated the contribution of [U-^13^C]glutamine, another major source for TCA cycle anaplerosis^43,44^ (Figure 1J). This revealed an even higher contribution of this amino acid as compared to glucose, highlighting glutamine as the primary TCA cycle carbon source in proliferating MuSCs and differentiated myotubes (Figure 1F, right). Upon differentiation to myotubes, the fractional contribution of glutamine to the TCA cycle decreased (Figure 1F), indicating an adaptation of the enzymatic network for glutamine utilization. In parallel to this global reprogramming, the directionality of contribution changed towards decreased reductive carboxylation over oxidative anaplerosis, as captured by the m+5 / m+4 ratio of citrate labeling (Figure 1K and S1E). Together, glutamine and glucose-derived carbons represented 71 % of TCA cycle carbons in myoblasts and 79 % in myotubes (Figure S1F), highlighting them as the primary sources feeding the TCA cycle. In conclusion, we observed a surge in TCA cycle contribution from glucose upon differentiation that was enabled by flux through PDH as well as anaplerosis via PC. This was paralleled by decreased utilization of glutamine as an anaplerotic substrate and a switch to more oxidative contributions of glutamine to the TCA cycle.

### Reprogrammed expression of metabolic genes enables TCA rewiring in myotubes

To elucidate the transcriptional changes underlying the metabolic reprogramming during differentiation, we next performed RNA sequencing in myoblasts and myotubes. Differentiation to myotubes was confirmed by characteristic marker gene expression including *Myogenin* and *Myosin heavy chain 3* and *4*, paralleled by a downregulation of *Pax7* and *Myod1* (Figure S2A). For unbiased identification of global expression changes, Gene Set Enrichment Analysis (GSEA) was performed using WikiPathways. The strongest enrichment of metabolic pathways in myotubes included electron transport chain and oxidative phosphorylation, glycolysis and gluconeogenesis, and TCA cycle (Figure 1L and S2B-S2D) while pathways associated with proliferation and cell cycle control were downregulated (Figure S2B). Moreover, *Pcx* (encoding PC) expression was upregulated in myotubes concomitant with a decrease in *Phosphoenolpyruvate Carboxykinase 2* (*Pck2*) (Figure S2E) mechanistically supporting the enhanced m+3 contribution of glucose to the TCA cycle via tracing. Focusing on potential transcriptionally controlled regulators of glutamine anaplerosis, all enzymes involved in conversion of glutamate to aKG by transamination were higher in myotubes, contrasting to the decreased contribution from glutamine. Only *Glud1*, a mitochondrial glutamate deaminase, was significantly decreased in myotubes (Figure 1M), suggesting that *Glud1* downregulation might contribute to a reduced glutamate to aKG conversion. The quantification of intracellular metabolites supported lower conversion, as evidenced by the increased glutamate / aKG ratio driven by decreased aKG concomitant with a higher abundance of glutamate and aspartate in myotubes (Figure 1D, 1N, and S1C). Importantly, the increase in glutamate occurred despite a reduction in *Glutaminase 1* (*Gls1*) expression (Figure S2F). GLS1 catalyzes the conversion of glutamine to glutamate which is the primary source of intracellular glutamate^45^. Along with the lower fractional contribution from GLS1 to the glutamate pool, glutamine-derived nitrogen contribution to glutamate, aspartate, and alanine was significantly decreased in myotubes (Figure S2G). Together, this indicates that myotubes use amino acids besides glutamine as nitrogen sources to generate glutamate and other non-essential amino acids from alpha-ketoacids. In conclusion, we found that glucose and glutamine metabolism rewire during myotube differentiation. Myotubes have a higher contribution of glucose to TCA cycle intermediates and lower glutamine anaplerosis when compared to myoblasts. Consistent with lower glutamine anaplerosis, we observed a decrease in the expression levels of the glutamine deaminase *Glud1* concomitant with an accumulation of glutamate, while transaminases using glutamate were upregulated. We thus wondered whether *Glud1* could play a role in MuSC differentiation.

### Loss of *Glud1* promotes myoblast differentiation leading to imbalanced myotube fusion *in vitro*

To study the role of GLUD1 in MuSCs, we next generated *Glud1* deficient satellite cells. *Pax7-CreER*^T2^ mice^46^ were intercrossed with *Glud1*^flox/flox^ mice^47^ to generate a *Pax7-CreER*^T2^-*Glud1*^flox/flox^ strain. We then isolated MuSCs from *Pax7-CreER*^T2^-*Glud1*^flox/flox^ mice, brought them in culture, and induced recombination *in vitro* by treating them with 1.5 μM tamoxifen for 5 consecutive days (*Glud1*^ΔSC^). Non-tamoxifen-treated cells were used as a control (WT), after confirming that tamoxifen treatment itself did not have any effect on differentiation (Figure 2A and S3A). Efficient recombination was confirmed using qPCR and Western blot (Figure 2B and 2C). Since preventing glutamine uptake in MuSC cell lines reduces proliferation ^38^, we first evaluated proliferation in WT and *Glud1*^ΔSC^ myoblasts. No difference was observed in the fraction of Ki67^+^ or EdU^+^ myoblasts between *Glud1*^ΔSC^ and WT myoblasts (Figure 2D and 2E), indicating a role of *Glud1* independent of fueling proliferation. Next, we assessed whether loss of *Glud1* affects MuSCs fate decisions. To do so, we determined the percentage of PAX3/7^+^ cells present in the myoblast cultures by flow cytometry. Interestingly, removing *Glud1* reduced the proportion of PAX3/7^+^ MuSCs when compared to WT (Figure 2F and S3B). Upon double staining for PAX7 and MYOD we observed that the decrease in PAX7^+^ cells in *Glud1*^ΔSC^ myoblast cultures was largely accounted for by a reduced proportion PAX7^+^ MYOD^-^ cells, (Figure 2G and S3C), showing that *Glud1*^ΔSC^ myoblasts lose their self-renewal capacity.

**Figure 2.**
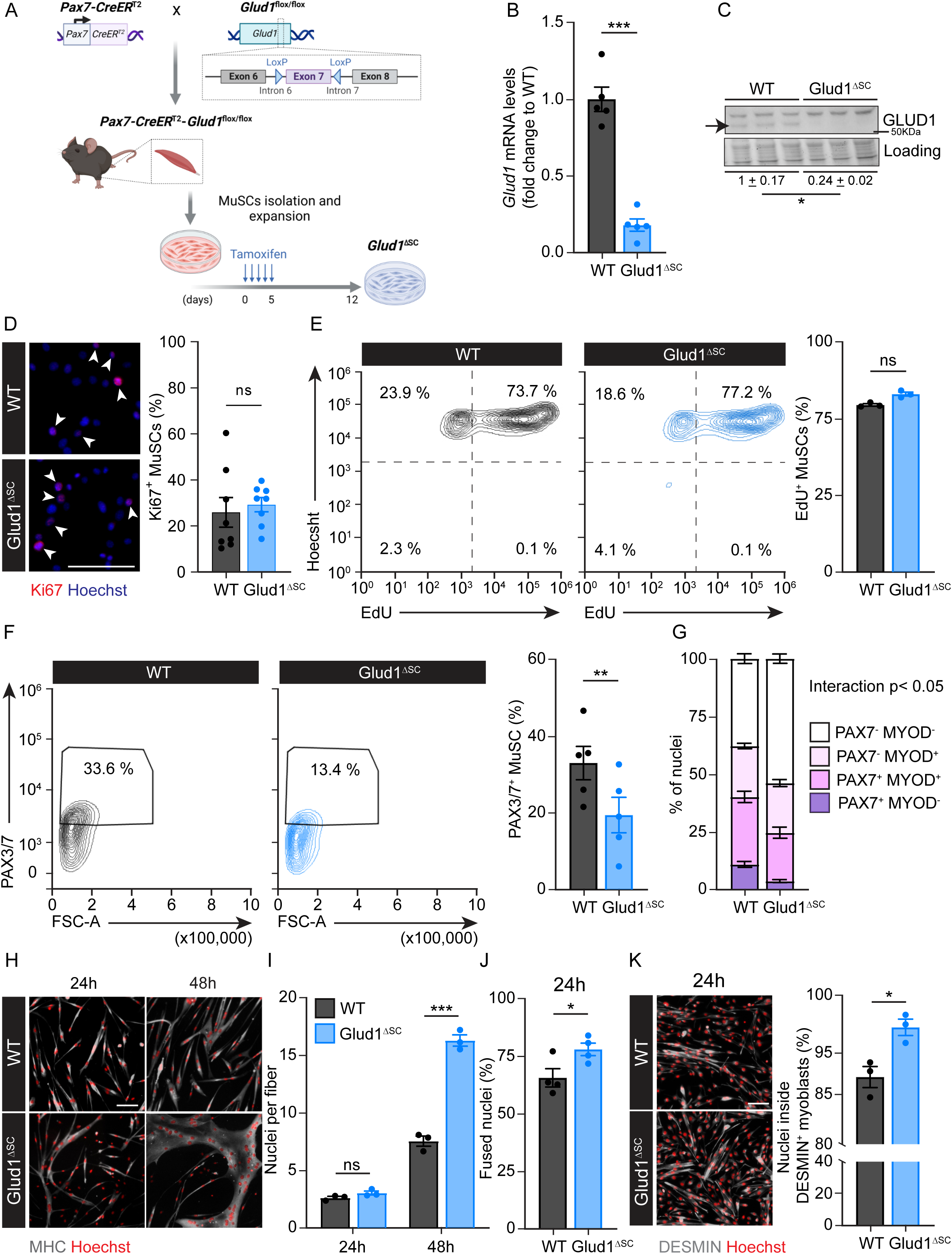
Loss of *Glud1* promotes myoblasts differentiation leading to imbalanced myotube fusion *in vitro*. (A) Schematic representation of the generation of *Pax7-CreER*^T2^x*Glud1*^flox/flox^ mice, experimental setup for MuSCs isolation, and tamoxifen treatment to generate *Glud1*^ΔSC^ MuSCs. (B) Gene expression analysis of *Glud1* in WT and *Glud1*^ΔSC^ myoblasts. Dots represent independent experiments. (C) Protein expression analysis of GLUD1 in WT and *Glud1*^ΔSC^ MuSCs. (D) Representative images (left) and quantification (right) of Ki67 immunofluorescent staining (red, Ki67; blue, Hoechst; scale bar 100 µm) in WT and Glud1^ΔSC^ myoblasts. Dots represent biological replicates. (E) Representative flow cytometric analysis (left) and quantification (right) of EdU^+^ WT and *Glud1*^ΔSC^ myoblasts. A representative experiment is shown. Dots represent biological replicates. (F) Representative flow cytometric analysis (left) and quantification (right) of Pax3/7^+^ WT and *Glud1*^ΔSC^ myoblasts. (G) Quantification of PAX7 and MYOD immunofluorescent staining in WT and *Glud1*^ΔSC^ myoblasts. A representative experiment of three experiments is shown. (H) Representative images of myosin heavy chain (MHC) immunofluorescent staining (grey, MHC; red, Hoechst; scale bar 100 µm) in WT and *Glud1*^ΔSC^ myotubes differentiated for 24 or 48 h. (I) Quantification of the number of nuclei per myotube identified as described in (H). Dots represent independent experiments. (J) Quantification of the fusion index (% of fused nuclei) in WT and *Glud1*^ΔSC^ myotubes 24 h after the induction of differentiation. Dots represent independent experiments. (K) Representative images (left) and quantification (right) of the percentage of nuclei within a DESMIN^+^ (DES) cell in a DES immunofluorescent staining (grey, desmin; red, hoechst; scale bar 100 µm) in WT and *Glud1*^ΔSC^ myotubes 24 h after the induction of differentiation. Dots represent independent experiments. Graph bars represent mean ± SEM. Student’s t test (two-tailed, unpaired) in (B, D-F, and J-K). Two-way ANOVA in (G) and Two-way ANOVA with Sidak’s multiple comparison test in (I). (*p < 0.05, **p < 0.01, ***p < 0.001, ****p<.0001)

We next evaluated whether *Glud1* deletion also affected later steps of myogenesis. To do so, WT and *Glud1*^ΔSC^ myoblasts were seeded in differentiation media to induce myogenic differentiation and followed until their fusion to myotubes. Cells were harvested 24 and 48 h after the start of differentiation to track the dynamics of myogenic progression over time. Interestingly, *Glud1*^ΔSC^ myoblasts differentiated faster and formed much larger myotubes than their WT counterparts (Figure 2H). While at 24 h after inducing differentiation the number of nuclei per fiber was not different between WT and *Glud1*^ΔSC^ cells (Figure 2H and 2I), both the fusion index (Figure 2J) and the proportion of nuclei inside DESMIN^+^ myotubes (Figure 2K) were increased, further supporting that *Glud1*^ΔSC^ myoblasts differentiate faster. By 48 h, these differences resulted in myotubes derived from *Glud1*^ΔSC^ myoblasts that contained more myonuclei than control myotubes indicating a hyperfusion phenotype (Figure 2H and 2I). Importantly, this was not secondary to a gradual loss of PAX7^+^ cells since inducing recombination immediately prior to differentiation, thereby starting with identical numbers of PAX7^+^ cell numbers, also promoted differentiation / fusion of *Glud1*^ΔSC^ myoblasts (Figure S3D). Thus, *Glud1* deletion limits the self-renewal capacity of myoblasts and promotes myoblast differentiation and fusion to myotubes.

### Loss of *Glud1* leads to premature MuSC fusion and loss of self-renewal upon voluntary running-wheel exercise

To study whether loss of *Glud1* also restricts MuSC self-renewal and promotes differentiation *in vivo*, we decided to perform MuSC-tracing experiments by generating a mouse model that allows genetic labelling of MuSCs and therefore the ability to track their incorporation into muscle fibers. To that end, we intercrossed *Pax7*-CreER^T2^-*Glud1*^flox/flox^ mice with *Rosa26*^mTmG/mTmG^ fate tracing mice^48^ to generate a *Pax7*-CreER^T2^-*Glud1^f^*^lox/flox^-*Rosa26*^mTmG/mTmG^ strain (*Glud1*^ΔSC-mG^). *Pax7*-*CreER*^T2^-*Glud1*^wt/wt^*-Rosa26*^mTmG/mTmG^ (*Glud1*^WT-mG^) offsprings were used as controls. In these mice tamoxifen induced CRE recombinase activation will lead to the excision of *mTomato* and expression of mGFP. This approach allows us to permanently label PAX7^+^ MuSCs, and thereby assess their self-renewal potential, as well as the incorporation of PAX7^+^ MuSCs into myofibers (Figure 3A). Because exercise is known to induce MuSC activation, proliferation and promotes their contribution to myofibers^6,8,13,27^, we next subjected *Glud1*^WT-mG^ and *Glud1*^ΔSC-mG^ mice to a voluntary resistance exercise protocol. Briefly, mice were exposed to 2 weeks of voluntary running wheel exercise without any resistance in the wheel, after which the resistance was gradually increased up to 65 % (week 3) and maintained at 65 % until the end of the exercise training (week 7) (Figure 3A). No differences in running distance were observed between genotypes (Figure 3B). Since *musculus (m.) soleus, m. plantaris* as well as *m. gastrocnemius* exhibit a significant increase in MuSC contribution to myofibers upon voluntary running wheel exercise ^27^, we used those muscles to determine the MuSC self-renewal capacity and differentiation upon *Glud1* loss. In the unperturbed state, we found approximately 80 % of mGFP^+^ PAX7^+^ MuSCs in both, *Glud1*^WT-mG^ and *Glud1*^ΔSC-mG^ mice, 1 week after tamoxifen treatment (Figure 3C and S3E). However, after training, the proportion of mGFP^+^ PAX7^+^ MuSCs in *Glud1*^ΔSC-mG^ mice (*m. plantaris* and *m. gastrocemius*) was reduced (Figure 3C and S3E) when compared to *Glud1*^WT-mG^ controls, showing impaired ability of MuSCs to self-renew and repopulate the niche after exercise-induced activation in *Glud1*^ΔSC-mG^ mice. We next evaluated whether *Glud1* deficient MuSCs also precociously differentiate, leading to reduced contribution to myofibers. In *Glud1*^WT-mG^ *m. soleus*, 30 % of fibers were mGFP^+^ after 7 weeks of running, indicating that at least 1 MuSC fused to each fiber (Figure 3D). In *Glud1*^ΔSC-mG^ the contribution from MuSCs to myofibers was significantly reduced, as indicated by lower mGFP^+^ fiber area after running (Figure 3D). Overall, these data suggest that *Glud1* deletion promotes imbalanced MuSC self-renewal and myotube fusion, both *in vitro* and *in vivo.* We subsequently wondered how GLUD1 controls MuSCs behavior.

**Figure 3.**
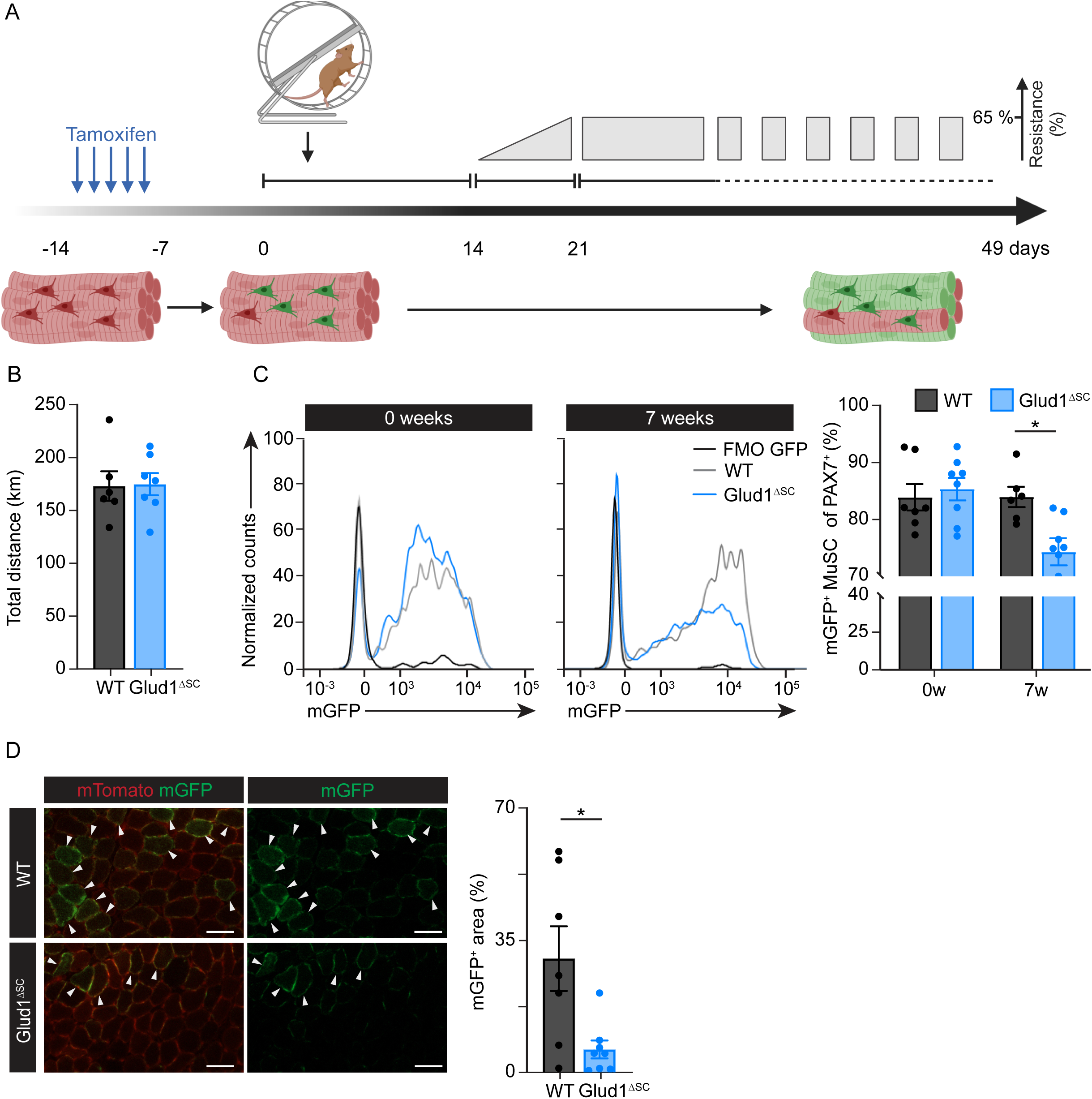
Loss of *Glud1* causes premature MuSC fusion and impaired self-renewal upon voluntary running-wheel exercise. (A) Schematic representation of the tamoxifen treatment schedule and the resistance running wheel protocol used to assess MuSC contribution to muscle fibers upon exercise. (B) Total distance run by WT and *Glud1*^ΔSC^ mice during the 65 % resistance phase. (C) Representative flow cytometric analysis (left) and quantification (right) of mGFP^+^ cells as a percentage of PAX7^+^ cells before (0w) and after (7w) the 7 week running wheel protocol. (D) Representative images (left) of endogenous mTomato and mGFP (red, mTomato; green, mGFP. Scale bar 50 µm) in WT and *Glud1*^ΔSC^ *m. soleus* muscle after 2 weeks of wheel running and quantification (right) of total mGFP^+^ area. White asterisks indicate mGFP^+^ myofibers. Graph bars represent mean ± SEM. Dots represent individual mice. (B and D) Student’s t-test (two tailed, unpaired) (C) Two-way ANOVA with Sidak’s multiple comparison test. (*p < 0.05, **p < 0.01, ***p < 0.001).

### Loss of *Glud1* does not affect the core transcriptional regulation factors of MuSC differentiation

To identify the molecular mechanisms leading to premature hyperfusion downstream of *Glud1* loss, we first analyzed the expression profile of myogenic genes by qPCR. Surprisingly, no significant differences in the expression levels of classic myogenic genes were observed between WT and *Glud1*^ΔSC^ myoblasts throughout myogenesis (Figure S3F). To further elucidate potential genetic changes in detail, we then used a previously described approach to uncouple myogenic differentiation from fusion^49,50^. Myoblasts were differentiated for 48 h at low density to prevent them from fusing. Then, the resulting myotubes were trypsinized and re-seeded at high confluency to allow fusion (Figure 4A). We then performed RNA sequencing on samples harvested in each of the conditions: proliferating myoblasts, fusion-competent myocytes, and fused multinuclear myotubes. Samples from both genotypes clustered together based on differentiation status, indicating that loss of *Glud1* leads to pronounced phenotypical changes while the transcriptomic landscape remained similar to WT throughout differentiation. (Figure 4B and 4C). This was highlighted by the fact that 94 % of the genes differentially expressed in WT myocytes as compared to myoblasts were in common with those differentially expressed in *Glud1*^ΔSC^ myocytes. Making the same comparisons between the myotube state and myoblast state, approximately 92 % of the genes differentially regulated in WT cells were shared with *Glud1*^ΔSC^ cells (Figure 4D). Finally, expression patterns of the main myogenic genes were equally changed over time (Figure 4E). Overall, loss of *Glud1* pushes myoblasts away from self-renewal towards differentiation causing imbalanced myotube fusion independent of myogenic gene expression. We therefore hypothesized that loss of *Glud1* metabolically primes myoblasts to differentiate faster.

**Figure 4.**
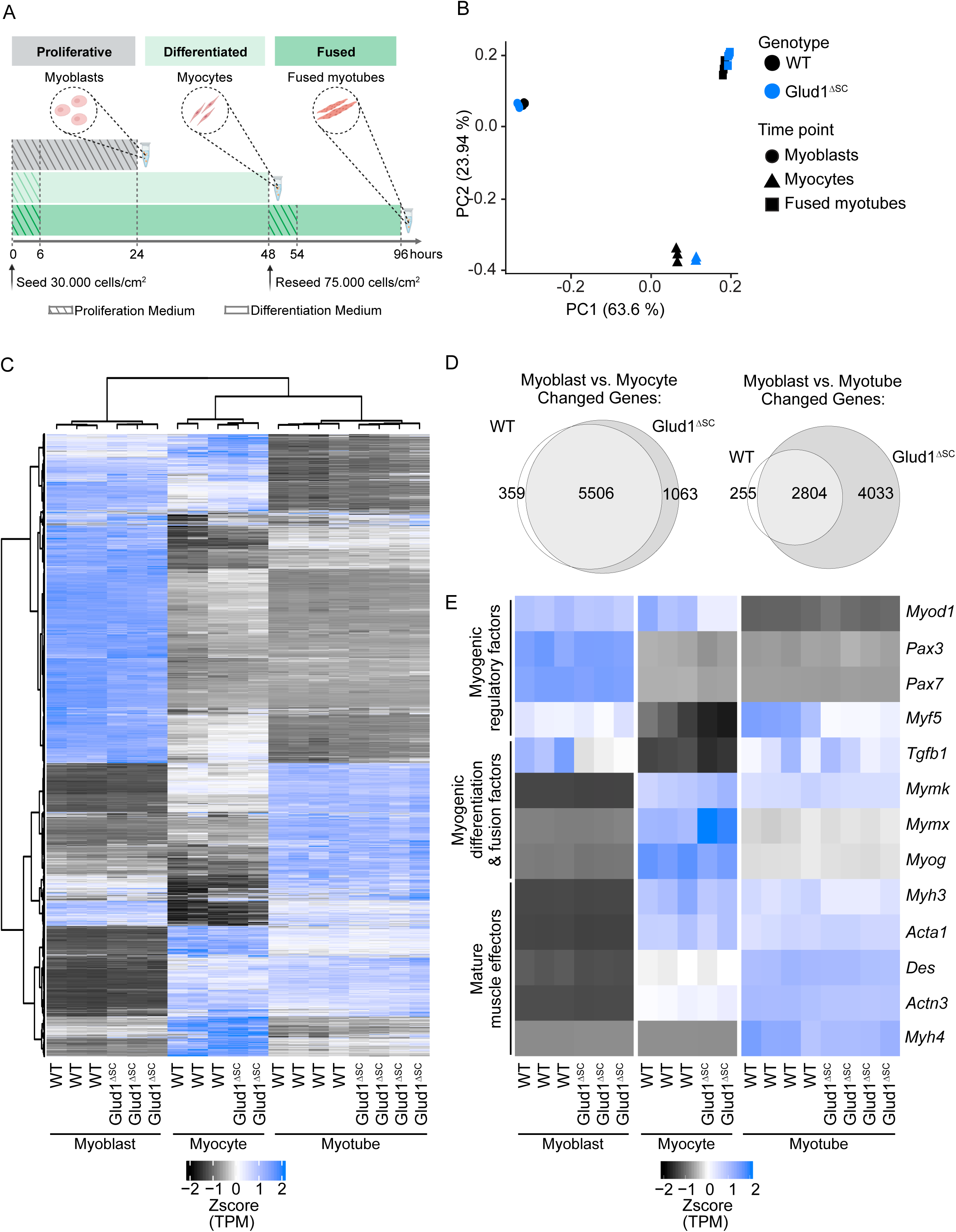
Loss of *Glud1* does not affect core transcriptional regulation of MuSC differentiation. (A) Schematic representation of the approach used to uncouple the processes of myogenic differentiation and fusion. (B) Principal component analysis of WT and *Glud1*^ΔSC^ cells at 3 different time points based on RNA-seq data. (C) Heatmap of expressed genes changed across conditions. Clustering was performed using the Ward method. (D) Euler diagram of changing genes, comparing myotube or myocyte to myoblast and evaluating overlap between genotypes. Changing genes were defined as genes with an absolute log2 fold change > 1 between conditions. (E) Expression of myogenic marker genes across conditions and genotypes. (B-E) Myoblast: n = 3 per genotype, Myocyte: WT n = 3 and *Glud1*^ΔSC^ n = 2, Myotube: n = 4 per genotype.

### Loss of *Glud1* leads to imbalanced nitrogen compartmentalization by glutamate trapping

To evaluate the metabolic changes associated with loss of *Glud1*, we performed metabolic analysis in cell lysates from WT and *Glud1*^ΔSC^ myoblasts. Surprisingly in [U-^13^C]glucose tracing the fractional contributions from glucose to the TCA cycle intermediates in *Glud1*^ΔSC^ myoblasts were unchanged when compared to WT (Figure 5A and S4A). Similarly, the contribution to TCA cycle intermediates from [U-^13^C]glutamine was unchanged between WT and *Glud1*^ΔSC^ myoblasts (Figure 5A and S4B). These data suggest that *Glud1* is not essential for glutamine anaplerosis to the TCA cycle and that the functional redundancy of glutamate dependent transaminases compensates for the exchange between glutamate and aKG in *Glud1*^ΔSC^ myoblasts. Moreover, when we evaluated the metabolite pool size of TCA intermediates and other metabolites directly related with glutamine metabolism such as glutamate, aspartate, alanine, and serine, we detected only increases in glutamate (Figure 5B) and fumarate (Figure S5A).

**Figure 5.**
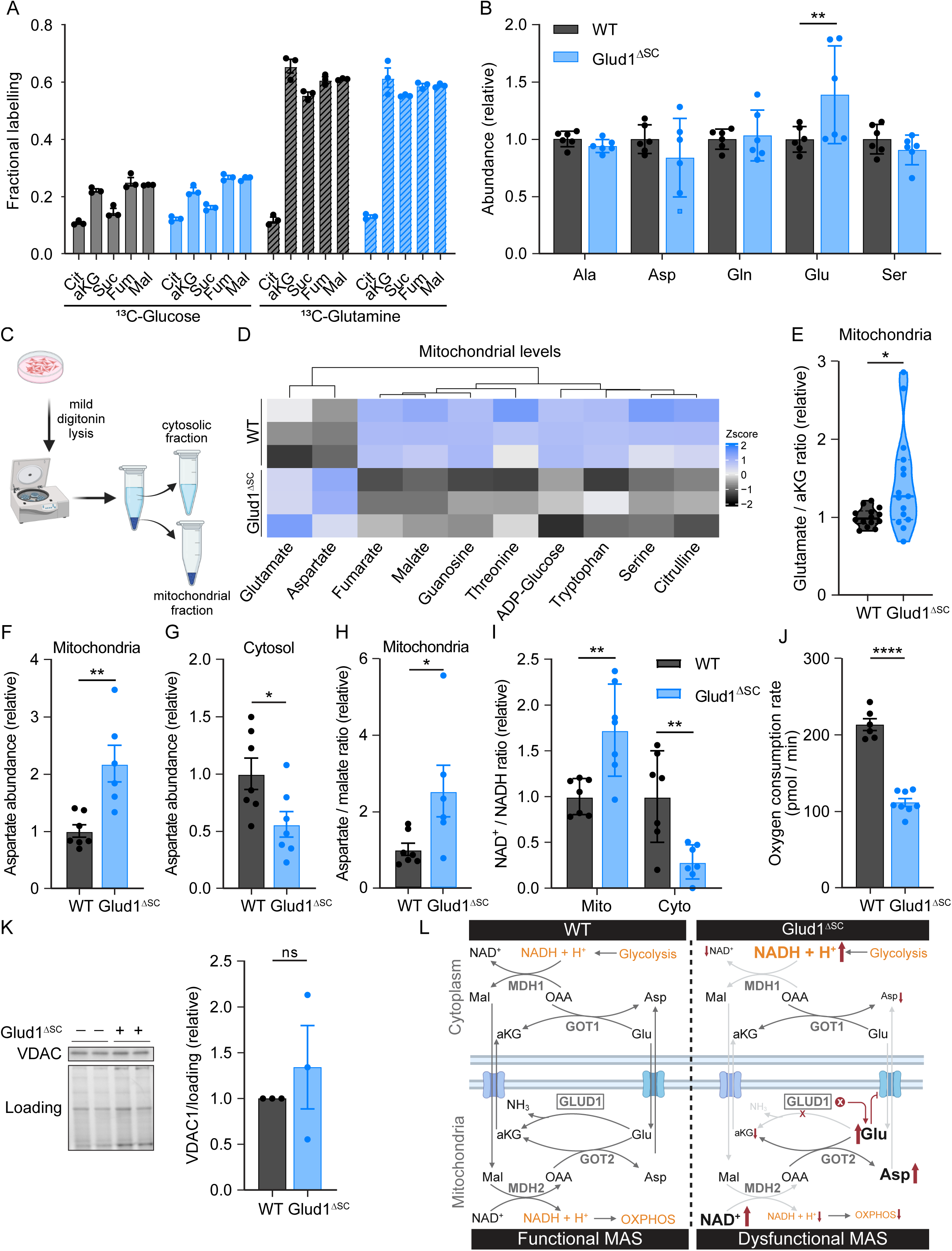
Loss of *Glud1* leads to imbalanced nitrogen compartmentalization. (A) Fractional contribution of [U-^13^C]glucose (left) or [U-^13^C]glutamine (right) to the TCA cycle metabolites citrate (Cit), alpha-ketoglutarate (aKG), succinate (Suc), fumarate (Fum), and malate (Mal) in WT and *Glud1*^ΔSC^ myoblasts. (B) Relative abundance of the amino acids alanine (Ala), aspartate (Asp), glutamine (Gln), glutamate (Glu), and serine (Ser) in WT and *Glud1*^Δ*SC*^ myoblasts. (C) Schematic representation of the subcellular fractionation protocol to isolate the mitochondrial and cytosolic fractions for metabolomics analysis. (D) Mitochondrial levels of top 10 most changed metabolites in WT compared to *Glud1*^ΔSC^ myoblasts. (E) Relative mitochondrial glutamate / aKG ratio in WT and *Glud1*^ΔSC^ myoblasts. (F) Relative mitochondrial aspartate levels in WT and *Glud1*^Δ*SC*^ myoblasts. (G) Relative cytosolic aspartate levels in WT and *Glud1*^Δ*SC*^ myoblasts. (H) Relative mitochondrial aspartate / malate ratio in WT and *Glud1*^Δ*SC*^ myoblasts. (I) Relative mitochondrial and cytosolic NAD^+^ / NADH ratio in WT and *Glud1*^Δ*SC*^ myoblasts. (J) Oxygen consumption rate in WT and *Glud1*^Δ*SC*^ myoblasts. (K) Representative pictures (left) and quantification (right) of western blot analysis of VDAC1 protein levels in WT and *Glud1*^Δ*SC*^ myoblasts. (L) Schematic representation of the flow of NADH into the mitochondria via the malate aspartate shuttle (MAS) and the dysfunction induced in *Glud1*^Δ*SC*^ myoblasts. Bar graphs represent mean ± SEM in (B) and (F-K). Bar graphs represent mean ± SEM in (A). Violin plot displays the mean (black line) and the quartiles (dashed line). Each dot represents a biological replicate in (A-B) and (F-J) or the average of an independent experiment in (K). Student’s t test (two-tailed, unpaired) in (E-H) and (J-K). Two-way ANOVA with Tukey’s multiple comparison test in (A). Two-way ANOVA with Sidak’s multiple comparison test in (I). (*p < 0.05, **p < 0.01, ***p < 0.001, ****p<.0001)

Because neither metabolite pool sizes nor fractional contribution of glutamine to the TCA cycle changed substantially after *Glud1* deletion (Figure 5A and 5B and S5A), we hypothesized that the role of GLUD1 could be restricted to a cellular compartment, rather than at the whole cell level. Indeed, GLUD1 is a deaminase that is only present in mitochondria, whereas most transaminases have mitochondrial and cytoplasmic isoforms (Figure S5B). Therefore, we set out to investigate whether loss of *Glud1* impacts metabolite compartmentalization. To do so, we performed subcellular fractioning followed by liquid chromatography-mass spectrometry (LC-MS) to determine metabolite abundance in both the mitochondrial and cytoplasmic fractions, as described previously^51,52^. The purity of the subcellular fractions was confirmed by western blot based on the abundance of VDAC1 and GAPDH, which mark the mitochondrial and cytosolic fractions, respectively (Figure S5C). Unlike in the whole cell lysates, we detected marked changes in metabolite abundance between WT and *Glud1*^ΔSC^ myoblasts in the mitochondrial fractions of many of the metabolites analyzed (Figure 5D). One of the most striking differences was the accumulation of glutamate relative to aKG (Figure 5E) as well as the accumulation of aspartate (Figure 5F), in the mitochondrial fractions of *Glud1*^ΔSC^ myoblasts. A concomitant decrease in aspartate was observed in the cytosolic fractions of *Glud1*^ΔSC^ myoblasts (Figure 5G), confirming that *Glud1* has a pivotal role in coordinating compartment-specific metabolite abundance. This compartment-specific metabolite imbalance led to an elevated mitochondrial aspartate / malate ratio (Figure 5H) and NAD^+^ / NADH ratio (Figure 5I), indicating a block in the malate-aspartate shuttle (MAS). The MAS serves as a net transport system of reducing equivalents from the cytosol to the mitochondrial respiratory chain and therefore feeds oxidative phosphorylation with NADH from glycolysis. The functional consequence was supported by a decrease in basal oxygen consumption rate (OCR) in *Glud1*^ΔSC^ myoblasts (Figure 5J) a direct redout of mitochondrial oxidative phosphorylation activity. The decrease in OCR was neither driven by changes in mitochondrial content (Figure 5K) nor transcriptional changes (Figure S5D), indicating that this decrease in OCR is likely an effect of the imbalanced mitochondrial NAD^+^ / NADH ratio rather than changes in the electron transport chain itself. Together our data indicate that loss of *Glud1* results in mitochondrial nitrogen accumulation in the form of glutamate. The glutamate accumulation results in a build-up of mitochondrial aspartate, due to insufficient nitrogen release in the form of ammonia, disrupting MAS activity (Figure 5L).

### Alanine supplementation rescues MAS deficiency and imbalanced fusion in *Glud1* deficient MuSCs

To elucidate the relevance of nitrogen compartmentalization on MAS function during myogenesis, we next tested whether amino acid supplementation could restore MAS activity, and therefore imbalanced fusion in *Glud1*^ΔSC^ myoblasts. Whereas glutamate and aspartate are poorly transported across the cellular membrane^53^, alanine is readily imported^53^, and acts as a cytosolic nitrogen donor via alanine aminotransferase. We therefore used alanine supplementation to contribute to the cytosolic nitrogen donor pool in myoblasts. WT and *Glud1*^ΔSC^ myoblasts were cultured in proliferation media supplemented with alanine for 24 h (Figure 6A). Metabolite abundance in the mitochondrial and cytoplasmic fractions was determined by LC-MS. Strikingly, alanine supplementation was sufficient to fully rescue the elevated mitochondrial aspartate / malate ratio (Figure 6B and S6A) and to increase cytosolic aspartate as well as glutamate levels in *Glud1*^ΔSC^ myoblasts (Figure 6C and 6D). Additionally, alanine supplementation in *Glud1*^ΔSC^ myoblasts resulted in full recovery of MAS shuttle activity, as evidenced by a normalization of both the cytosolic and mitochondrial NAD^+^ / NADH ratios to levels observed in WT myoblasts (Figure 6E and S6B). This indicates that equilibrating nitrogen compartmentalization on the aspartate-glutamate axis by supplementing alanine is enough to fully rescue the MAS dysfunction despite persistent deletion of *Glud1*.

**Figure 6.**
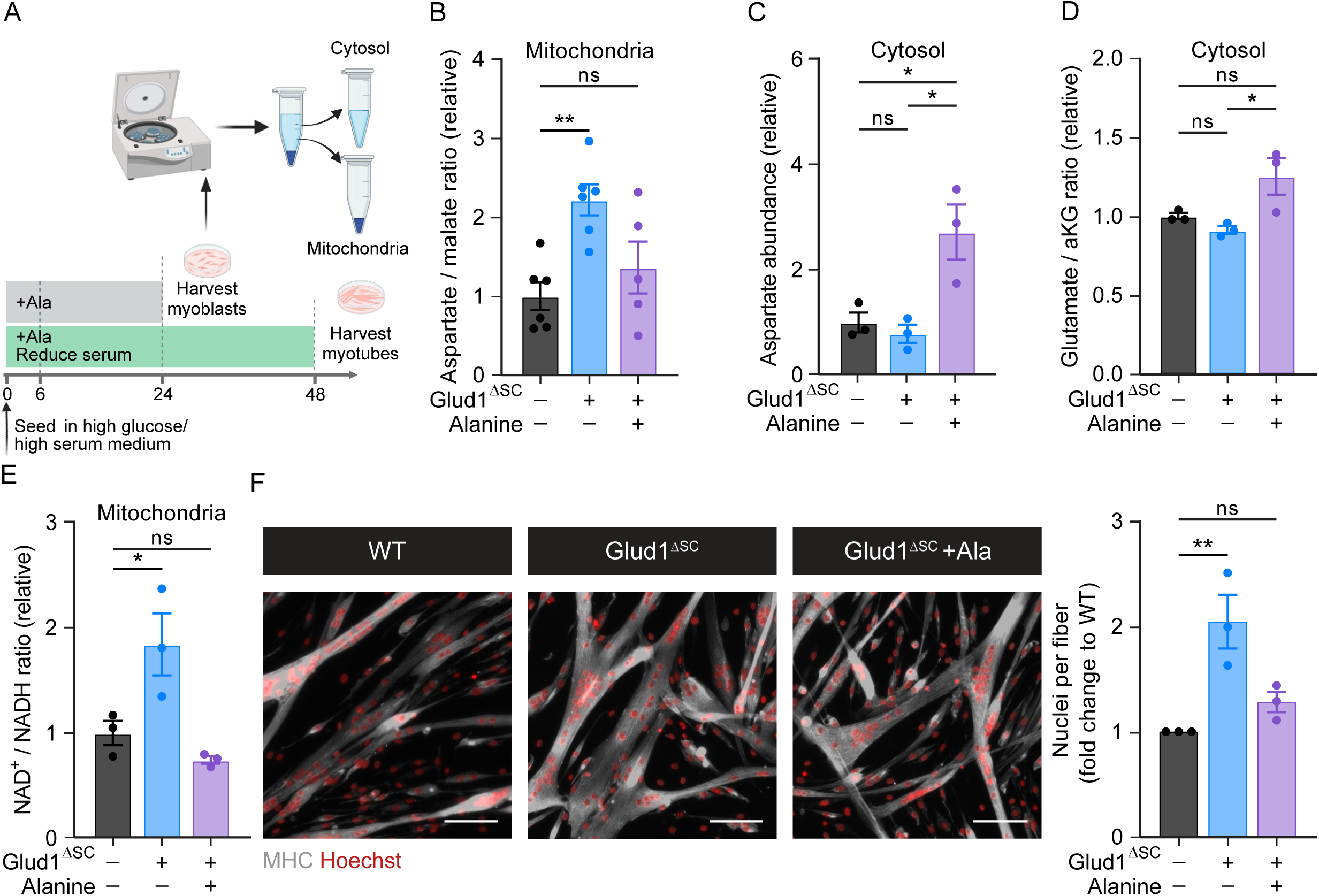
Alanine supplementation rescues MAS deficiency and imbalanced fusion in *Glud1* deficient MuSCs. (A) Schematic representation of the alanine supplementation protocol for metabolomics at the myoblast stage (top) and differentiation analysis at the myotube stage (bottom). (B) Relative mitochondrial aspartate / malate ratio in WT and *Glud1*^ΔSC^ myoblasts grown in growth media with or without 0.2 mM alanine supplementation. (C) Relative cytosolic aspartate levels in WT and *Glud1*^ΔSC^ myoblasts grown in growth media with or without 0.2 mM alanine supplementation. (D) Relative cytosolic glutamate / alpha-ketoglutarate ratio in WT and *Glud1*^ΔSC^ myoblasts grown in growth media with or without 0.2 mM alanine supplementation. (E) Relative mitochondrial NAD^+^ / NADH levels in WT and *Glud1*^ΔSC^ myoblasts grown in growth media with or without 0.2 mM alanine supplementation. (F) Representative images of myosin heavy chain (MHC) immunofluorescent staining (grey, MYHC; red, hoechst, scale bar 100 µm) in WT and *Glud1*^ΔSC^ myotubes differentiated in media with or without 0.2 mM alanine supplementation (left) and quantification (right) of the nuclei per fiber relative to WT without alanine supplementation. Bar graphs represent mean ± SEM in (B-E). Each dot represents a biological replicate in (B-E) or the average of an independent experiment in (F). One-way ANOVA with Dunnet’s multiple comparison test in (B and E). One-way ANOVA with Tukey’s multiple comparison test in (C and D) (*p < 0.05, **p < 0.01, ***p < 0.001, ****p<.0001).

We then asked whether the metabolic phenotype rescue was paralleled by normalization of *Glud1*^ΔSC^-induced imbalanced fusion. In this respect, myoblast fusion was assessed in WT and *Glud1^ΔSC^*myoblasts after culture in alanine supplemented differentiation media for 48 h (Figure 6A). In line with the restoration of the metabolic phenotype, alanine supplementation normalized fusion in *Glud1*^ΔSC^ myotubes to WT levels (Figure 6F), confirming the direct link between myogenic differentiation and compartmentalized nitrogen balance. Overall, these data show that supplementation of alanine restores MAS activity in *Glud1* deficient MuSCs and prevents normalizes metabolic function and imbalanced fusion. We thereby highlight a crucial contribution of compartmentalized amino acid metabolism for MuSC fate decisions.

## Discussion

Recent work has shown the existence of non-canonical TCA cycling in proliferating embryonic stem cells as well as a skeletal muscle stem cell line (C2C12)^33^. Compared to proliferating C2C12 myoblasts, differentiated myotubes increased glucose incorporation into the TCA cycle and switched from non-canonical to canonical glucose TCA cycle activity. At least partially, this switch was mediated via the activation of pyruvate dehydrogenase^31,33^. Consistently, we found increased glucose contribution to the TCA cycle via pyruvate dehydrogenase upon differentiation, as indicated by enhanced m+2 labeling of TCA cycle intermediates. In addition, myotubes activated pyruvate carboxylase dependent anaplerosis from glucose, a feature typically observed for glutamine dependent cells upon glutamine withdrawal^54^, suggesting active regulation of glutamine anaplerosis during differentiation. Indeed, similar to more committed embryonic stem cells^55^ and many mammalian cell lines^43,44,56^, we found that proliferating MuSCs use glutamine as their dominant TCA cycle substrate. The glutamine contribution to the TCA cycle was both oxidative, as well as reductive, suggesting that it supports the generation of citrate, which can be used for non-canonical cycling.

The importance of glutamine uptake and metabolism for MuSC function is still poorly understood. Previous work has shown that depletion of intracellular glutamine in MuSCs by preventing glutamine availability or uptake impairs MuSC proliferation and differentiation^37^. These effects were dependent on mTORC1, a known regulator of MuSC activation and proliferation^32,57^. We did not observe a proliferation defect upon loss of G*lud1,* potentially because glutamine availability and total intracellular glutamine levels were not affected by *Glud1* deletion. Also, deleting *Glud1* from MuSCs did not affect the contribution of glutamine to the TCA cycle, likely because transamination was able to compensate. While both glutamine and glutamate are crucial metabolites within the glutamine anaplerotic pathway, it thus seems that they control MuSC fate via different mechanisms. In support of an alternative mechanism, inhibition of glutamine uptake or reducing glutamine availability reduced *Myogenin* expression^37^ and inhibition of mTORC1 reduced the expression of several myogenic genes^57^. In contrast, upon deletion of *Glud1,* we neither observed major transcriptional reprogramming nor differences in pathways specifically associated with myoblast differentiation and fusion.

Instead, we found that when glutamine is available, *Glud1* controls glutamate abundance in the mitochondria. Active *Glud1* in myoblasts thereby keeps mitochondrial glutamate levels in check to allow efficient TCA cycle anaplerosis during proliferation without a build-up of mitochondrial glutamate that would induce precocious differentiation. By doing so it allows proliferating myoblasts to be greedy with glutamine without being pushed towards differentiation. Genetic deletion of *Glud1* resulted in compartment specific accumulation of glutamate and pushed muscle stem cells towards differentiation and imbalanced fusion. At the same time, it reduced the self-renewal capacity of MuSCs as evidenced by the rapid reduction in the number of PAX7^+^ cells. However, this lower self-renewal capacity did not suffice to explain the increased differentiation / fusion, since inducing differentiation immediately after tamoxifen treatment showed an unchanged phenotype. The phenotypic presentation occurred in the absence of a transcriptional switch and was dependent on the activity of the MAS. Restoring MAS deficiency, induced by *Glud1* deletion through alanine supplementation completely rescued the precocious differentiation and fusion. Our data highlights that metabolism, in addition to the known genetic regulators, can directly control MuSC fate and fusion phenotype.

The MAS transports reducing equivalents across the mitochondrial membrane. This transmembrane shuttle thereby allows the provision of reducing equivalents from glycolysis, generated in the form of NADH, for energy production by the electron transport chain (ETC). Concomitantly it regenerates cytosolic NAD^+^ to keep a favorable NAD^+^ / NADH ratio for glycolytic flux and pathways such as serine biosynthesis. This net transport into the mitochondria is critically dependent on the coupled export of mitochondrial aspartate with an import of cytosolic glutamate followed by an exchange of cytosolic malate for mitochondrial aKG. In MuSCs, *Glud1* deletion induced a build-up of mitochondrial glutamate which caused MAS deficiency evidenced by accumulation of cytosolic NADH along with accumulation of mitochondrial NAD^+^ and lower oxygen consumption by the ETC. This build-up of mitochondrial glutamate is opposite to expected changes upon direct MAS inhibition by impairment of the mitochondrial glutamate / aspartate transporter (SLC25A12). Opposed to our findings this would be expected to induce depletion of mitochondrial glutamate as SLC25A12 facilitates the import of glutamate into the mitochondria coupled with aspartate export. No gross histological changes have been observed in the muscle of SLC25A12 knock-out mice nor humans with SLC25A12 deficiency^58–60^, whereas the effect of these impairments on MuSC fate decisions *in vivo* has not been investigated. *In vitro* experiments have shown though that SLC25A12 is crucial for cycling MuSCs, which rely on glutamate-aspartate exchange between the mitochondrial and the cytosolic compartments^61^. Knock down of SLC25A12 lead to decreased proliferation in C2C12 myoblasts, whereas we did not observe a difference in proliferation in *Glud1* ^ΔSC^ myoblasts. This could indicate that while a complete block of the MAS inhibits proliferation, more subtle activity changes by trapping in the mitochondria, as induced by the glutamate build-up in *Glud1*^ΔSC^ myoblasts, is not sufficient to decrease MuSC proliferation.

Exercise is a known activator of MuSCs in mice and humans^13–15^, leading to their contribution to myofibers in a load-dependent manner. Upon muscle stem cell activation, they undergo rounds of proliferation to expand the stem cell pool. The majority of those cells differentiate to myocytes, while a smaller portion undergoes asymmetric division to self-renew. *In vivo*, the loss of self-renewal and precocious differentiation of activated MuSCs reduced their contribution to muscle fibers and depleted the muscle stem cell pool. Thus, active mitochondrial GLUD1 is required for proper *in vivo* stem cell contribution to the active muscle fibers.

In summary, we find that mitochondrial glutamate balance regulated by GLUD1 is critical to retain MuSCs in an undifferentiated state and prohibits their differentiation and fusion. Losing this break causes mitochondrial buildup of glutamate and aspartate inducing a dysfunctional malate / aspartate shuttle.

## Supporting information

Soro-Arnaiz et al_Supplementary Information

## Acknowledgements

We thank the De Bock and Morscher groups as well as Junyoung O. Park for scientific discussions and insights. We thank the Functional Genomics Center Zürich (FGCZ) for excellent technical support. I.S.A. was supported by the Fondation Suisse de Recherche sur les maladies musculaires. K.D.B. is endowed by the Schulthess Foundation. G.F., S.C. and R.J.M. were supported by grants from the NOMIS Foundation, Holcim-Stiftung Wissen, Gertrud-Hagmann-Stiftung für Malignom-Forschung, EMDO-Stiftung and Heidi Ras Grant of the FZK University Children’s Hospital Zürich. J.C.G.C. is supported by a grant from the Conselleria de Sanidad Universal y Salud Pública, Generalitat Valenciana, as part of Plan GenT, Generació Talent (DEI-01/20-C). All the schematic representations included in this paper were created with BioRender.com

## Author contributions

I.S.A. conceptualized the study, designed, performed, and analyzed experiments, and wrote the manuscript. S.C. and G.F. designed, performed and analyzed experiments and wrote the paper. J.Z., P.G. and E.M., contributed to the performance of experiments and data analysis. A.G. performed bioinformatics analysis under the supervision of O.B.N.. P.M. provided the Glud1 floxed mice. N.Z, M.P., A.C. and J.C.G.C performed metabolomics measurements. K.D.B. and R.J.M. conceptualized the study, supervised the experiments and data analysis, acquired funding and wrote the paper.

## STAR METHODS

### RESOURCE AVAILABILITY

#### Lead contact

Further information and requests for resources and reagents should be directed to and will be fulfilled by the Lead Contacts, Katrien De Bock and Raphael J. Morscher.

#### Materials availability

This study did not generate new unique reagents.

#### Data and code availability

The RNA-seq data reported in this study are available at the Gene Expression Omnibus (GEO) repository under the accession number (GSE 243640). All other data are available from the corresponding authors on request.

### EXPERIMENTAL MODELS

#### Mice

All mouse lines were maintained on a C57BL/6J background: Pax7-CreER^T2^ (The Jackson Laboratory strain# 017763), Rosa26^mTmG/mTmG^ (The Jackson Laboratory strain# 007576), and Glud1^flox/flox^ (*Glud1^tm^*^1^*^.1Pma^*, MGI:3835667)^47^ mice. To obtain inducible MuSC-specific G*lud1-* knockout mice, Glud1^flox/flox^ mice were intercrossed with Pax7-CreER^T2^ mice to generate Pax7-CreER^T2^-*Glud1*^flox/flox^ mice. To perform MuSCs-tracing experiments *in vivo*, Pax7-CreER^T2^-*Glud1*^flox/flox^ mice were intercrossed with Rosa26^mTmG/mTmG^ mice to generate *Pax7*-Cre^ERT2^-*Glud1^flox/flox^*-*Rosa26^mTmG/mTmG^* mice. *Pax7*-Cre^ERT2^-*Glud1*^wt/wt^-*Rosa26^mTmG/mTmG^* mice were used as controls. At the age of 8-10 weeks, mice received tamoxifen via intraperitoneal injection at a dose of 1.0 mg / day for 5 days to induce CRE-mediated recombination, and a washout period of 7 days was allowed before experiments were started.

Mice were randomly allocated to different experimental groups, and the investigator was blinded to the group allocation during the experiment as well as during the analysis. Mice were housed in individually ventilated cages at standard housing conditions (22 °C, 12 h light / dark cycle, dark phase starting at 7 pm), with *ad libitum* access to chow diet (18 % proteins, 4.5 % fibers, 4.5 % fat, 6.3 % ashes, Provimi Kliba SA) and water. Health status of all mouse lines was regularly monitored according to FELASA guidelines. All animal experiments were approved by the local animal ethics committee (Kantonales Veterinärsamt Zürich, licenses ZH180/18, and ZH039/21), and performed according to local guidelines (TschV, Zurich) and the Swiss animal protection law (TschG).

##### Exercise-induced MuSC activation

Mice were individually housed in open cages equipped with a running wheel device (TSE Systems, Bad Homburg vor der Höhe, Germany) for the duration of the intervention. A representation of the study design is shown in Figure 4A. Prior to the 4 weeks of voluntary wheel running, mice were familiarized for 14 days to the running wheel without extra resistance. After familiarization, mice were subjected to a progressive increase in resistance on the wheel: The load on the wheel was 50 % from day 14 to 17, 60 % from day 18 to 20, 65 % from day 21 until the end of the experiment. All mice ran at 65 % resistance during the last 4 weeks. Mice were euthanized and hind-limb muscles were harvested. *M. soleus* was frozen in OCT embedding matrix (CellPath) in liquid nitrogen-cooled isopentane for histological analysis, and *m. gastrocnemius* and *m. plantaris* were harvested and processed for MuSC isolation for flow cytometry (see below).

#### Cell culture

##### Standard culture conditions

Isolated primary mouse MuSCs were cultured in proliferation media (1 : 1 ratio of low-glucose DMEM (Cat#22320-022, ThermoFisher Scientific) and Ham’s F-10 (1 X) nutrient mix (Cat#22390-025, ThermoFisher Scientific) supplemented with 30 % FBS (Cat#10270106, Thermo Scientific) heat inactivated at 56 °C for 30 min and 10 ng / ml human basic-FGF (Cat#PHG0266, ThermoFisher Scientific) on dishes coated with Matrigel Basement Membrane Matrix (Cat#356237, Corning, 1 : 15 dilution). To induce myogenic differentiation MuSCs were seeded at 30.000 cells / cm^2^ on plates coated with Matrigel and cultured in differentiation medium: high-glucose DMEM, Cat#11995073, ThermoFisher Scientific) supplemented with 0.2 % FBS-heat inactivated (Cat#10500064, ThermoFisher Scientific) for 2 days.

##### Culture media for metabolomic analysis

For metabolic characterization, proliferation and differentiation media were adapted to exclude the influence of the media in the analysis. Proliferation media contained DMEM high glucose supplemented with 30 % dialyzed FBS (Cat#26400044, ThermoFisher Scientific) and 10 ng / ml basic-FGF. Before plating the experiment, MuSCs were cultured for 2 passages (maximum 3 days) in proliferation media to ensure adaptation. Differentiation media for metabolomic analysis contained DMEM high glucose supplemented with 0.2 % dialyzed FBS and 29.8 % PBS (Cat#10 0-015, ThermoFisher Scientific), indicating that only the growth factor content was reduced. MuSCs were seeded at 30.000 cells / cm^2^ on plates coated with Matrigel in proliferation media for metabolomic analysis. For the myoblasts condition, MuSCs were cultured in proliferation media for 24 h and then harvested for metabolite extraction (see below). For the myotube condition, after 24 h in proliferation media, MuSCs were washed once with PBS and differentiation was induced by culturing the cells with differentiation media. After 48 h, myotubes were harvested for metabolite extraction (see below)

All media were supplemented with 100 units / ml penicillin and 100 μg / ml streptomycin (Cat#15140122, ThermoFisher Scientific). Cells were routinely cultured at 37° C in 21 % O_2_ and 5 % CO_2_. Cells were regularly tested for the presence of mycoplasma. Isolation was performed from male and female mice and similar results were obtained.

##### In vitro tamoxifen treatment

primary MuSCs were treated with 1.5 µm Tamoxifen (Cat#T5648, Sigma-Aldrich) for 5 consecutive days. After a wash-out period of 3-5 days, cells were used to plate experiments. For acute tamoxifen treatment (Figure S2B), 1.5 µm Tamoxifen was added at the same time as the differentiation media was added (induction of differentiation). Tamoxifen was kept in the media until the end of the experiment (48 h).

##### Uncoupled differentiation-fusion protocol

This protocol was adapted from *Girardi et al., 2021*^49^. Briefly, primary murine myoblasts were seeded at a low density (5000 cells / cm^2^) in proliferation media to avoid contact between cells and therefore fusion. Six hours after seeding, the media was changed to differentiation media to induce differentiation. Cells were kept in differentiation media for 48 h. The resulting cultures after this step are called differentiated fusion competent myocytes throughout the manuscript. Then, differentiated fusion competent myocytes were trypsinized (Cat#25200056, ThermoFisher Scientific) and re-seeded at high confluency (75 000 cells / cm^2^) in proliferation media. Six hours after seeding, the media was changed to differentiation media to allow myotube fusion and kept in culture for 48 h. The resulting cultures after this step are called fused myotubes throughout the paper. (See Figure 4A).

##### Alanine supplementation

For alanine (Cat#A7469-100, Sigma-Aldrich) supplementation experiments, alanine was supplemented 6 h after seeding to reach a final concentration of 0.3 mM in the media, as opposed to the 0.1 mM basal alanine concentration.

### METHOD DETAILS

#### Isolation of mouse MuSC

For primary mouse MuSC isolation, mice were euthanized, all hind-limb muscles were immediately dissected, and muscles were minced in a petri dish on ice using a surgical blade. Next, the minced muscle tissue was enzymatically digested in digestion buffer containing 0.2 % Collagenase II (Cat#17101015, ThermoFisher Scientific), and 1.5 % BSA (Cat#A1391.0100, AppliChem) in HBSS (Cat#14025100, ThermoFisher Scientific) at 37 °C for 40 min with constant shaking. The reaction was stopped by adding an equal volume of TC media containing low glucose (1 g / L) DMEM supplemented with 10 % FBS, and the suspension was passed through a 100 μm cell strainer (Cat#FAL352360, Falcon). Subsequently the suspension was centrifuged at 350 g for 5 min at 4 °C and the supernatant discarded. The pellet was then suspended in 2 ml of erythrocyte lysis buffer (154 mM NH_4_CL, 10 mM KHCO_3_, 0.1 mM EDTA, pH = 7.35) and incubated on ice for approximately 1 min (longer for samples with more blood). 20 mL of cold PBS was then added to the suspension which was then 50 g for 5 min. After centrifugation, the supernatant was collected and immediately filtered through a 40 μm cell strainer (Cat#FAL352360, Falcon) to remove tissue debris. Cell suspension was centrifuged at 650 g for 8 min at 4 °C. The pellet was then suspended in 3 % BSA in PBS and centrifuged at 550 g for 5min. This step was repeated at least 3 times to wash the cell suspension. After the last centrifugation, the cell pellet was suspended in MuSCs proliferation media and plated on a non-coated plate. Cell suspension was pre-plated at 37 °C in 21 % O_2_ and 5 % CO_2_ for 3 h to remove the fibroblasts from cell preparation. While fibroblasts quickly attach to non-coated plates, MuSCs are unable to do so. Therefore, after 3 h in the incubator most fibroblasts will stay attached on the uncoated plate and MuSCs can be recovered by collecting the media from the plate. Then the MuSCs-enriched suspension was plated to a Matrigel-coated plate and kept in culture for MuSCs expansion. The pre-plating step was repeated at least 3 times during the next passages. MuSCs between passage 6 and 18 were used to perform experiments in this study.

#### Metabolomic characterization

##### Metabolite extraction

Cells were grown in 6 cm dishes. Myoblasts were harvested 24 h after seeding and myotubes were harvested 48 h after induction of differentiation (Figure 1A). After removing the media, cells were washed once with 2 ml of ice-cold PBS and 1 ml of quenching solution was added immediately 40:40:20 (v / v / v) acetonitrile:methanol:water. The quenching solution was pre-cooled at -80 °C for 2 h). Plates were immediately scraped on ice and the lysates were transferred to a pre-cooled 1.5 ml eppendorf tube. After a centrifugation (16626.896 g, 5 min at 4 °C), 800 µm of the supernatant were collected in a clean pre-cooled 1.5 ml eppendorf tube and stored at -80 °C until analysis.

##### Glucose and glutamine tracing experiments

The culture conditions for tracing experiments were the same as explained above for the regular metabolomic analysis (see cell culture section) with the difference that 8 h prior to cell harvesting proliferation or differentiation media was replaced by fresh media containing stable isotope-labelled glucose or glutamine. (Cat#CLM-1396-2, CLM-1822, and NLM-1328; Cambridge Isotope Laboratories).

#### Rapid mitochondrial fractionation

Rapid mitochondrial fractionation was performed as previously described^52^. In detail, cells were grown in 6 cm dishes and harvested 24 h after seeding in the myoblast state. After removing the media, the cells were washed twice with 2 ml of ice-cold PBS. Subsequently, cells were scraped from the cultured dishes and collected into a 1.5 mL tube using 1 ml of ice-cold PBS. After rapid centrifugation (13,500 g, 10 s, 4 °C), the supernatant was removed and the cell pellet was resuspended in freshly prepared ice-cold digitonin (Cat#D141, Sigma-Aldrich) solution (1 mg / ml digitonin in PBS). The cell pellet was triturated 5 times with a p1000 pipette. The sample was then centrifuged (13,500 g, 10 s, 4 °C). After centrifugation, the supernatant was transferred to another 1.5 ml tube as the cytosolic fraction and the pellet remains as the mitochondrial fraction. For western blotting assays, the mitochondrial fraction was resuspended in 1 ml ice cold PBS. Both the mitochondrial and cytosolic fractions were then sonicated (3 x 10 s, 4 °C). For each sample, 333 µL of 4 x Laemmli Buffer was added, the sample was boiled for 5 min, and then loaded onto the gel. For metabolomics analysis, the mitochondrial fraction was immediately quenched by the addition of -80°C-chilled 50:30:20 (v / v / v) methanol:acetonitrile:water. The cytosolic fraction was quenched by the addition of 4 ml -80°C-chilled 62.5:37.5 (v / v) methanol:acetonitrile.

#### Metabolomics

##### Reverse phase chromatography method

MS preparation and processing was performed as previously published^62^. Briefly, extracts were lyophilized overnight and resolubilized in loading buffer according to PCV (water and 0.5 % formic acid) in narrow-bottom 96-deep-well plates on a shaker (800 rpm., 15 °C, 10 min) for LC–MS injection. Metabolites were separated using an Ultimate 3000-RS (Thermo Fisher Scientific) equipped with an ACQUITY UPLC HSS T3 1.8-µm, 100 × 2.1 mm internal diameter column (Waters) and eluted using the following gradient from solvent A (water, 5 mM ammonium formate and 0.1 % formic acid) to solvent B (methanol, 5 mM ammonium formate and 0.1 % formic acid) as follows: 2 min at 0 % B; 2–3.5 min to 4 % B; 3.5–10 min to 45 % B; 10–12 min to 70 % B; 12–13.5 min to 100 % B; with an isocratic plateau at 100 % B for 2–15.5 min and from 15.5–16.5 min to 0 % B. After each run the column was re-equilibrated for 8 min at 100 % A with a constant flow rate of 0.4 ml min^−1^. Mass spectra were acquired using a heated ESI source of a Q-Exactive high-resolution mass spectrometer (Thermo Fisher Scientific). Mass spectra were recorded in positive and negative mode with the MS detector in full-scan mode (full MS) in a scan range 50–750 m / z with an AGC target of 1 × 10^6^, a resolution of 70,000 and a maximum injection time of 80 ms. Heated ESI parameters were sheath gas flow rate 35 arbitrary units (AU), auxiliary gas flow rate 35 AU, sweep gas flow rate 2 AU, spray voltage 3.5 kV, capillary temperature 350 °C and aux gas heater temperature 350 °C. Detector settings for full MS were in-source CID 0.0 eV; µscans of 1; resolution of 70,000; AGC target of 1 × 106; max IT of 35 ms and spectrum data type, profile.

##### Hydrophilic interaction chromatography method

Clean extracts were dried down in a SpeedVac Vacuum Concentrator (Thermo Fischer Scientific) and resolubilized in a suitable volume of loading buffer (40:40:20, methanol:acetonitrile:water) according to PCV. LC separation was done on a XBridge BEH Amide column (2.1 mm × 150 mm, 2.5 μm particle size; Waters) using a gradient of solvent A (20 mM ammonium acetate, 20 mM ammonium hydroxide in 95:5 water:acetonitrile, pH 9.45) and solvent B (acetonitrile). The flow rate was 150 μL / min, column temperature was 25 °C, autosampler temperature was 5 °C, and injection volume was 10 μL. The LC gradient was as follows: 0 min, 90 % B; 2 min, 85 % B; 3 min, 75 % B; 7 min, 75 % B; 8 min, 70 % B; 9 min, 70 % B; 10 min, 50 % B; 12 min, 50 % B; 13 min, 25 % B; 14 min, 25 % B; 16 min, 0 % B; 21 min, 0 % B; 22 min, 90 % B; 25 min, 90 % B. The autosampler temperature was 5 °C, and the injection volume was 10 μL. The mass spectrometer was operated in negative ion mode to scan from m / z 70 to 700 at 1 Hz, a resolving power of 140,000, AGC target 3e6, and maximum injection time, 200 ms; and from m / z 650 to 1200 at 1 Hz and a resolving power of 140,000, AGC target 5e5, and maximum injection time, 500 ms. Data was acquired in centroid mode. Other MS parameters are as follows: sheath gas flow rate, 30 (arbitrary units); aux gas flow rate, 5 (arbitrary units); sweep gas flow rate, 0 (arbitrary units); spray voltage, 3.4 kV; capillary temperature, 320 °C; S-lens RF level, 50.

##### Data Analysis

LC-MS raw data files (.raw) were converted to mzXML format using msconvert (3.0.20315, ProteoWizard). Peaks were integrated with El-Maven (v0.0.12.1, Elucidata) using windows of 10 ppm and retention time as previously determined using a library of standards. For tracer experiments, isotope labeling was corrected for 13C and 15N natural abundances using the AccuCor package^63^. For the metabolic pathway enrichment analysis, the KEGG database was parsed using the KEGGREST package (Bioconductor, 3.17) to extract all metabolites from each pathway. Metabolites were ranked according to their significance (unpaired two-tailed Student t-test) between MB and MT and hypergeometric tests were performed to compute pathway enrichment, taking into account the metabolome coverage of the experiment.

#### Absolute quantification of amino acids

The concentrations of 50 amino acids in the sample were measured in an accredited laboratory using a validated analytical method. Briefly, to 50 µL of sample volume 50 µL of a solution containing 10 % sulfosalicylic acid and 50 isotopically labelled amino acids were added. The sample was thoroughly mixed and centrifuged at 4156.724 g for 10 min to precipitate proteins. To 10 µL of the resulting supernatant 70 µL borate buffer and 20 µL of a supersaturated AccQ·Tag solution (6-aminoquinolyl-N-hydroxysuccinimidylcarbamate) were added, which allows the derivatization of all primary and secondary amino groups. Derivatization was carried out by incubating the sample at 55 °C for 10 min. Thereafter, the sample was cooled to room temperature for 15 min. After dilution, samples were transferred to an autosampler set to 10 °C. Chromatographic separation was achieved using a CORTECS UPLC C18 reverse-phase column (2.1×150 mm, 1.6 µm particle size and 90 Å pore size, Waters) kept at 55 °C using an ACQUITY H-Class UPLC system (Waters) interfaced to an ACQUITY QDa mass spectrometer (Waters). Mobile phase A and B consisted of 0.1 % formic acid in water and 0.1 % formic acid in acetonitrile, respectively. The flow rate was set to 500 µL / min and the injection volume was 10 µL. The gradient consisted of 99 % A (0 – 1 min), 99 % A to 87 % A (1 – 4 min), 87 % A to 85 % A (4 – 8.5 min), 85 % A to 5 % A (8.5 – 9.5 min), 5 % A (9.5 – 10.5 min), 5 % A to 99 % A (10.5 – 10.7 min), 99 % A (10.7 – 12 min). Ions were measured in positive mode and cone voltage was set to 10 V for all compounds. Data analysis was performed using the MassLynx software (Waters). Quantification was done using calibrators containing amino acids at known concentrations (Waters), which were analyzed at the beginning of each sample batch. At the beginning and at the end of each batch two internal quality controls were measured for quality control purposes.

#### Oxygen consumption (OCR)

Oxygen consumption was determined using a Seahorse XF-96 Extracellular Flux Analyzer according to the providers instructions (Seahorse Bioscience). Myoblasts were seeded at 50,000 cells / well in matrigel-coated 96-well plates. The assay medium was regular proliferation media. The measurements were performed for 5 cycles as follows: 30 s mixing, 2 min recovery, 3 min measuring.

#### RNA extraction and quantitative RT-PCR

RNA of myoblasts and myotubes was extracted using a RNeasy Plus Micro Kit according to the manufacturer’s instructions (Cat#74034, QIAGEN). RNA purity and concentration were assessed via a spectrophotometer (Tecan, Spark). RNA was reverse-transcribed to cDNA by High Capacity cDNA Reverse Transcription Kit (Cat#43-688-13, ThermoFisher Scientific). A SYBR Green-based master mix (Cat# A25778, ThermoFisher Scientific) was used for real-time qPCR analysis with primers listed in Table S2. To compensate for variations in RNA input and efficiency of reverse-transcription, 18S was used as a housekeeping gene. The delta-delta C_T_ method was used to normalize the data.

#### RNA sequencing and differential gene expression analysis

##### Library preparation

The quality of the isolated RNA was determined with a Fragment Analyzer (Agilent, Santa Clara, California, USA). The TruSeq Stranded mRNA (Illumina, Inc, California, USA) was used in the succeeding steps. Briefly, total RNA samples (100-1000 ng) were poly A enriched and then reverse-transcribed into double-stranded cDNA. The cDNA samples were fragmented, end-repaired and adenylated before ligation of TruSeq adapters containing unique dual indices (UDI) for multiplexing. Fragments containing TruSeq adapters on both ends were selectively enriched with PCR. The quality and quantity of the enriched libraries were validated using the Fragment Analyzer (Agilent, Santa Clara, California, USA). The product is a smear with an average fragment size of approximately 260 bp. The libraries were normalized to 10 nM in Tris-Cl 10 mM, pH8.5 with 0.1 % Tween 20.

##### Cluster Generation and Sequencing

The Novaseq 6000 (Illumina, Inc, California, USA) was used for cluster generation and sequencing according to standard protocols. Sequencing configuration was single-end 100 bp.

##### Analysis

Raw sequencing data was processed using the SUSHI framework^64,65^ developed at the Functional Genomics Center Zurich (FGCZ). Quality control was performed by trimming adapter sequences and low-quality reads using fastp v0.20^64^. Pseudo-alignment of the filtered reads was performed against the mouse reference genome assembly GRCm38.p6 and gene expression level was quantified using Kallisto v0.46.1^66^. Differential gene expression analysis was performed between conditions using the R package edgeR v3.42^67^. Gene set enrichment was performed using the leading-edge analysis^68^, using a gene list ranked by relative changes (log2 fold-change) as input. Gene sets were taken from the WikiPathways subset of the MSigDB C2 curated gene sets^69^. We used 1,000 permutations of the gene-levels values to calculate normalized enrichment scores and statistical significance. For visualization of the whole transcriptome, genes with sufficient coverage were identified and displayed using the ‘filterByExpr’ function from the R package edgeR v3.42^67^.

#### Immunoblot analysis

Cells were collected and lysed with RIPA buffer [50 mM Tris-HCl pH 7.4, 150 mM NaCl, 2 mM EDTA, 1.0 % Triton X-100, 0.5 % sodium deoxycholate] supplemented with 0.2 % SDS and cOmplete™ Protease Inhibitor Cocktail (Cat#11697498001, Roche). Lysates were centrifuged at 10,000 g for 10 min at 4 °C. Supernatant was collected, and protein concentration was measured using the DC protein assay kit (Cat#5000116, Bio-rad). 10-15 µg of total protein was loaded in a 4–15 % Mini-PROTEAN pre-casted gradient gel (Cat#456-8086, Bio-rad). After electrophoresis, a picture of the gel was taken under UV-light to determine total protein loading using stain-free technology. Proteins were transferred onto a PVDF membrane (1620177, Bio-rad) with a semi-dry system and subsequently blocked for 1 h at room temperature with 5 % milk in 0.1 % TBS-Tween. Membranes were incubated overnight at 4 °C with the following primary antibodies: anti-GLUD1 (Cat# ab34786, Abcam) and anti-VDAC1 (Cat#ab15895, Abcam). The appropriate HRP-linked secondary antibodies (rabbit IgG, HRP-linked Antibody, Cat#7074S, Cell Signaling Technology) were used for chemiluminescent detection of proteins. Membranes were scanned with a Chemidoc imaging system (Bio-rad) and quantified using Image Lab 6 software (Bio-rad). Total protein loading quantification was used for correction of gel loading differences.

#### Immunohistochemistry and histology

##### In vivo studies

*M. Soleus* muscle samples were harvested and embedded in Tissue-Tek and frozen in liquid N_2_-cooled isopentane. Frozen sections (10 μm) of muscle embedded in OCT of mid-belly level were made using a cryostat (Leica CM 1950) and collected on Superfrost Ultra Plus slides (10417002, ThermoFisher Scientific). MuSC contribution to muscle fibers in Pax7-CreER^T2^-Rosa26^mTmG/mTmG^ was quantified based on the % of mGFP positive area in the whole muscle section using ImageJ software. Images from skeletal muscle cryosections were captured at 10x using an epifluorescent microscope (Zeiss Axio observer Z.1, Zeiss, Oberkochen, Germany). Composite images were stitched together using the tiles module in the ZEN 2011 imaging software (Zeiss). All images were captured at the same exposure time. *<u>Myoblasts commitment and self-renewal</u>*: Myoblasts were fixed with 4 % PFA at room temperature for 7 min and washed twice with PBS. After permeabilizing for 1 min with PBS supplemented with 0.5 % Triton, blocking solution was added and cells were incubated for 1 h with blocking solution (2 % BSA in PBS supplemented with 0.1 % Triton X-100). Then, cells were incubated over-night at 4 °C with mouse IgG1 anti-Pax7 (MAB1675, R&D Systems, 5 μg / ml) diluted 1:100 in blocking solution and rat anti-MyoD (Cat#39991, Active Motif) diluted 1:100 in blocking solution. Then, cells were washed twice in blocking solution and incubated with goat anti-mouse IgG1, Alexa Fluor 568 (Cat#A-21124, ThermoFisher Scientific) diluted 1:250 in blocking solution and donkey anti-rat IgG, Alexa Fluor 488 (Cat#A-21208, ThermoFisher Scientific) diluted 1:250 in blocking solution. After washing twice with PBS, nuclei were counter stained with Hoechst (1:5000) (Cat#H3570, ThermoFisher Scientic). The staining solution was then removed, and cells were washed 2 times with PBS. Cells were imaged using AxioObserver.Z1 fluorescence microscope (Carl Zeiss, Oberkochen, Germany).

##### Differentiation and fusion analysis in vitro

Myoblasts or myotubes were fixed with 4 % PFA at room temperature for 7 min and washed twice with PBS. After blocking for 30 min with blocking solution (2 % BSA in PBS supplemented with 0.1 % Triton X-100), cells were incubated over-night at 4 °C with the following primary antibodies: Mouse monoclonal IgG2B (kappa light chain) anti-MHC (MF 20, DSHB), diluted 1:50 in blocking solution; or rabbit IgG anti-Desmin (Cat# 5332, Cell Signaling Technology), diluted 1:100 in blocking solution. Then, cells were washed twice in blocking solution and the incubate with the following secondary antibodies for 1 h at room temperature: Goat anti-mouse IgG2B, Alexa Fluor 488 conjugate (Cat#A21141, ThermoFisher Scientific), diluted 1:400 in blocking solution; goat anti-rabbit IgG, Alexa Fluor 568 conjugate (Cat#A11011, ThermoFisher Scientific), diluted 1:400 in blocking solution. After washing twice with PBS, nuclei were counter stained with Hoechst (1:5000). The staining solution was then removed and cells were washed 2 times with PBS. Cells were imaged using AxioObserver.Z1 fluorescence microscope (Carl Zeiss, Oberkochen, Germany).

##### Proliferation assays based on Ki67 staining

Myoblasts were fixed with 4 % PFA at room temperature for 7 min and washed twice with PBS. After permeabilizing for 1 min with PBS + 0.5 % Triton, blocking solution was added and cells were incubated for 1 h with blocking solution (2 % BSA in PBS supplemented with 0.1% Triton X-100). Then, cells were incubated over-night at 4 °C with rabbit anti-Ki67 (Cat#ab15580, AbCam) diluted 1:100 in blocking solution. Then, cells were washed twice in blocking solution and incubated with donkey anti-rabbit IgG, Alexa Fluor 488 (Cat#A21206, ThermoFisher Scientific), diluted 1:250 in blocking solution. After washing twice with PBS, nuclei were counter stained with Hoechst (1:5000). The staining solution was then removed and cells were washed 2 times with PBS. Cells were imaged using AxioObserver.Z1 fluorescence microscope (Carl Zeiss, Oberkochen, Germany).

#### Flow cytometry

##### In vivo MuSC self-renewal

For *in vivo* flow cytometry assay a protocol adapted from *Machado et al, 2017*^70^ was used. Briefly, mice were euthanized, *m. gastrocnemius* and *m. plataris* muscles were immediately dissected, and muscles were minced on a petri dish with 0.5 % PFA on ice using a surgical blade. Next, the minced muscle tissue was fixed for 1 h at 4 °C with constant shaking, washed twice with PBS and enzymatically digested in digestion buffer containing 2 mg / ml Collagenase IV (Cat#17104019, ThermoFisher Scientific), 2 mg / ml Dispase II (Cat#04 942 078 001, Roche Diagnostics), and 1.5 % BSA in HBSS at 37 °C for 45-60 min with constant shaking. The reaction was stopped by adding an equal volume of ice-cold low glucose (1 g / L) DMEM supplemented with 10 % FBS, and the suspension was passed through a 100 μm cell strainer. Subsequently the suspension was centrifuged at 600 g for 5 min at 4 °C and the supernatant discarded. The pellet was then suspended in ice-cold low-glucose DMEM, filtered through a 70 µm cell strainer (FAL352350, Falcon) and spun down for 5 min at 50 g at 4 °C. Then, the supernatant was filtered through a 40 µm cell strainer and spun down at 600 g for 5 min. Thereafter, cell pellets were kept and used for analysis after incubated with the appropriate primary antibodies. First, surface marker staining was performed by incubating the cell suspension for 1 h at 4 °C with the following antibodies: Brilliant Violet 605™ anti-mouse CD31 Antibody (Cat#102427, BioLegend, 1:100), Brilliant Violet 605™ anti-mouse CD45 Antibody (Cat#103139, BioLegend, 1:200), and Brilliant Violet 605™ anti-mouse Ly-6A/E (Sca-1) Antibody (Cat#108133, BioLegend, 1:100). Thereafter, samples were washed and permeabilized using BD Pharmingen™ Transcription Factor Buffer Set following manufacturer’s instructions (Cat#562574, BD Phamigen), followed by intracellular staining. To do so, cell suspensions were incubated with anti-PAX3/7, Alexa Fluor® 647 (B-5) (Cat#sc-365843 AF647, Santa Cruz Biothechnology) for 1 h at 4 °C diluted in 3 % BSA. Last, cells were counterstained with Hoechst and washed twice prior to analysis. Cells were analysed with a LSRFortessa (BD Bioscience) flow cytometer and data were analyzed using FlowJo 10 software (Tree Star).

##### In vitro PAX3/7 staining

Myoblasts were trypsinized and washed once with PBS. Fixation and permeabilization steps were performed using BD Pharmingen™ Transcription Factor Buffer Set following manufacturer’s instructions. For PAX3/7 staining, myoblasts were incubated with anti-PAX3/7, Alexa Fluor® 647 (B-5) (Cat#sc-365843 AF647, Santa Cruz Biothechnology) for 1 h at 4 °C diluted in 3 % BSA. Thereafter myoblasts were washed twice in 3 % BSA in PBS, counterstained with Hoechst and washed twice prior to analysis. Cells were analyzed on an SH800S sorter (Sony Biotechnology) and data were analyzed using FlowJo 10 software (Tree Star).

##### In vitro EdU incorporation assay

To assess entry into the cell cycle, incorporation of 5-ethynyl-2’-deoxyuridine (EdU, Cat#E10187, ThermoFisher Scientific) during the last 6 h in culture was assessed using the Click-iT Cell Reaction Buffer Kit (Cat#C10269, ThermoFisher Scientific) according to the manufacturer’s instructions. Briefly, myoblasts were incubated for 6 h in the presence of 10 µm EdU. Thereafter, the cells were trypsinized, washed once with PBS, and fixed with 4 % PFA for 15 min at 4 °C, and finally washed twice with 3 % BSA in PBS. After fixation, cells were permeabilized for 20 min at room temperature in 0.5 % Triton X-100 with 3 % BSA in PBS, then washed twice with 3 % BSA in PBS and incubated with the Click-iT reaction cocktail for 45 min in the dark at room temperature. Thereafter, cells were briefly washed and counterstained with Hoechst. Cells were analyzed on an SH800S sorter (Sony Biotechnology). Data were analyzed using FlowJo 10 software (Tree Star).

### QUANTIFICATION AND STATISTICAL ANALYSIS

The images presented in the manuscript are representative of the data and the image / staining quality. All data represent mean ± SEM. GraphPhad Prism software (version 8.0.0) was used for statistical analyses. Investigators were always blinded to group allocation. When comparing two group means, Student’s *t*-test was used in an unpaired two-tailed fashion. For more than two groups, one-way ANOVA with Tukey’s multiple comparisons test was used and for experimental set-ups with a second variable, two-way ANOVA with Sidak’s multiple comparisons test was used. The statistical method used for each experiment is indicated in each figure legend. Asterisks in figure legends denote statistical significance (*p < 0.05, **p < 0.01, ***p < 0.001). No experiment-wide multiple test correction was applied.

