## Supplementary material for "GLUD1 dictates muscle stem cell differentiation by controlling mitochondrial glutamate levels": Soro-Arnaiz et al_Supplementary Information

**Figure S1**

A

| Growth medium | | Differentiation medium | |
|---|---|---|---|
| DMEM High Glucose | 70 % | DMEM High Glucose | 70 % |
| dFBS | 30 % | dFBS | 0.2 % |
| PBS | - | PBS | 29.8 % |
| bFGF | 10 ng / ml | bFGF | - |
| Pen / Strep | 1 % | Pen / Strep | 1 % |

B

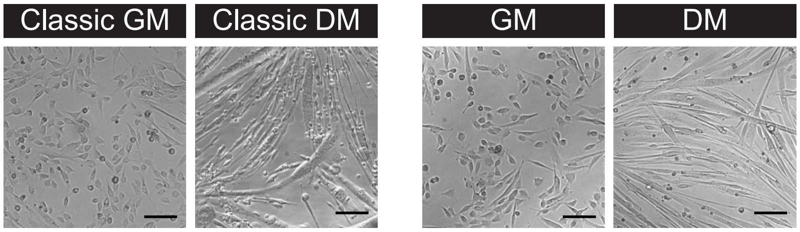


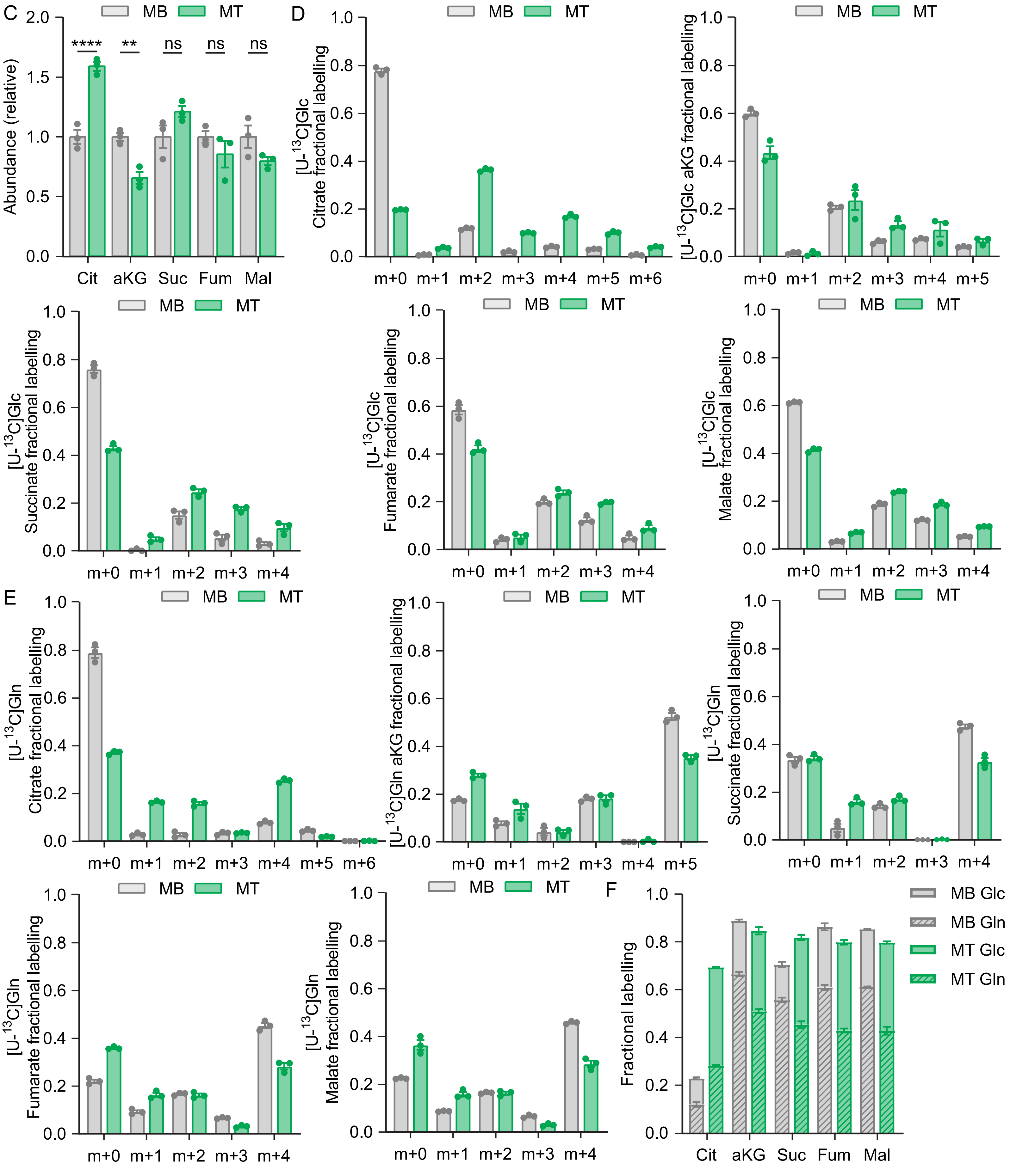

**Figure S1. A differentiation protocol allowing metabolomics studies highlights metabolic reprogramming consistent with gene expression changes. Related to Figure 1.**

(A) Table showing the main components of the growth and differentiation media used for growth and differentiation followed by metabolomic analysis including DMEM high glucose (4.5 g / L), dialyzed fetal bovine serum (dFBS), phosphate buffered saline (PBS), basic human fibroblast growth factor (bFGF) and penicillin / streptomycin (Pen / Strep). (B) Brightfield images of myoblasts grown in classic growth medium (classic GM) or in high glucose high dialyzed serum growth medium (GM) and myotubes differentiated in classic differentiation medium (classic DM) or in high glucose low dialyzed serum medium (DM). Scale bar 100 μm. (C) Relative abundance of the TCA cycle intermediates citrate (Cit), alpha-ketoglutarate (aKG), succinate (Suc), fumarate (Fum), and malate (Mal) in myoblasts (MB) and myotubes (MT). (D) Mass distribution vector (MDV) of citrate, alpha-ketoglutarate (aKG), succinate, fumarate and malate during [U-$^{13}$C]Glc tracing in MB and MT. (E) MDV of citrate, aKG, succinate, fumarate and malate during [U-$^{13}$C]Gln tracing in MB and MT. (F) Total fractional labelling of the TCA cycle intermediates citrate (Cit), alpha-ketoglutarate (aKG), succinate (Suc), fumarate (Fum), and malate (Mal) from [U-$^{13}$C]glucose and [U-$^{13}$C]glutamine in MB and MT. Bar graphs represent mean ± SEM. Each dot represents a biological replicate. Two-way ANOVA with Sidak's multiple comparison test in (C). (*$p < 0.05$, **$p < 0.01$, ***$p < 0.001$, ****$p<.0001$).

Figure S2

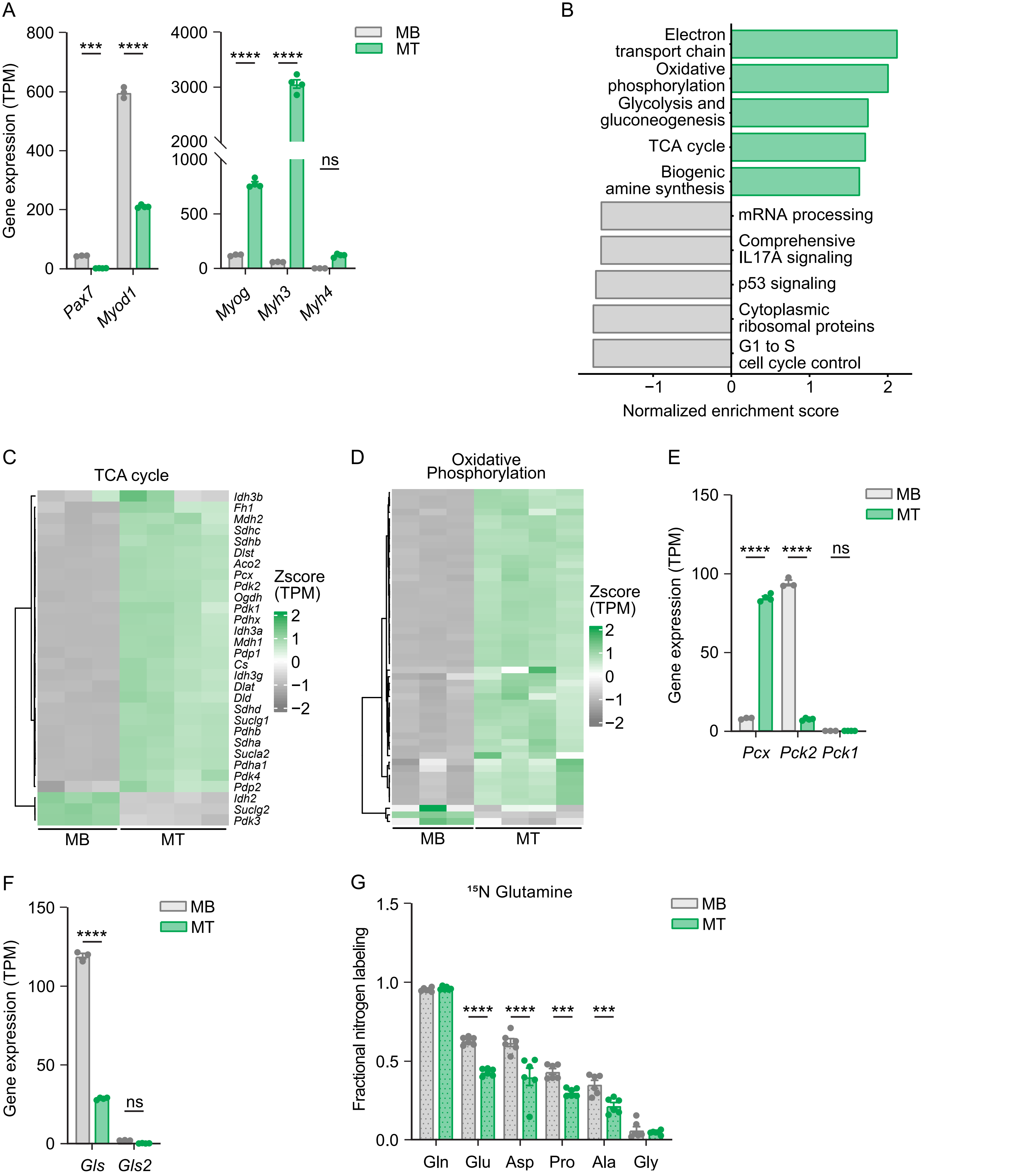

A
Gene expression (TPM)
Pax7
Myod1
Myog
Myh3
Myh4
MB
MT
ns
B
Electron transport chain
Oxidative phosphorylation
Glycolysis and gluconeogenesis
TCA cycle
Biogenic amine synthesis
mRNA processing
Comprehensive IL17A signaling
p53 signaling
Cytoplasmic ribosomal proteins
G1 to S cell cycle control
Normalized enrichment score
C
TCA cycle
Idh3b
Fh1
Mdh2
Sdhc
Sdhb
Dlst
Aco2
Pcx
Pdk2
Ogdh
Pdk1
Pdhx
Idh3a
Mdh1
Pdp1
Cs
Idh3g
Dlat
Dld
Sdhd
Suclg1
Pdhb
Sdha
Sucla2
Pdha1
Pdk4
Pdp2
Idh2
Suclg2
Pdk3
Zscore (TPM)
MB
MT
D
Oxidative Phosphorylation
Zscore (TPM)
E
Gene expression (TPM)
Pcx
Pck2
Pck1
F
Gene expression (TPM)
Gls
Gls2
G
¹⁵N Glutamine
Fractional nitrogen labeling
Gln
Glu
Asp
Pro
Ala
Gly

**Figure S2. Gene expression changes highlight metabolic reprogramming in muscle stem cell differentiation. Related to Figure 1.**

(A) Gene expression analysis of the canonical myogenic genes *Pax7, Myod1, Myog, Myh3, and Myh4* in MB and MT. (B) Significantly changing pathways (FDR < 0.05) identified using gene set enrichment analysis of WikiPathways. Positive normalized enrichment score (NES) denotes higher expression in MT (in green) and negative NES denotes higher expression in MB (in grey) (C) Expression of TCA cycle genes in MB and MT. Expression is evaluated using transcript per million (TPM) z-scored for each gene (D) Expression of oxidative phosphorylation genes in MB and MT. Expression is evaluated using TPM z-scored for each gene. (E) Gene expression analysis of *Pcx* (encoding PC)*, Pck2,* and *Pck1* in MB and MT*.* (F) Gene expression of the glutaminases *Gls1* and *Gls2* in MB and MT (G) Total fractional contribution of [$^{15}$N]glutamine to the nitrogen of the amino acids glutamine (Gln), glutamate (Glu), aspartate (Asp), proline (Pro), alanine (Ala), and glycine (Gly) in MB and MT. Bar graphs represent mean $\pm$ SEM. Each dot represents a biological replicate. Two-way ANOVA with Sidak's multiple comparison test in (C-E), (G), (K-M). (*p < 0.05, **p < 0.01, ***p < 0.001, ****p<.0001).

### Figure S3

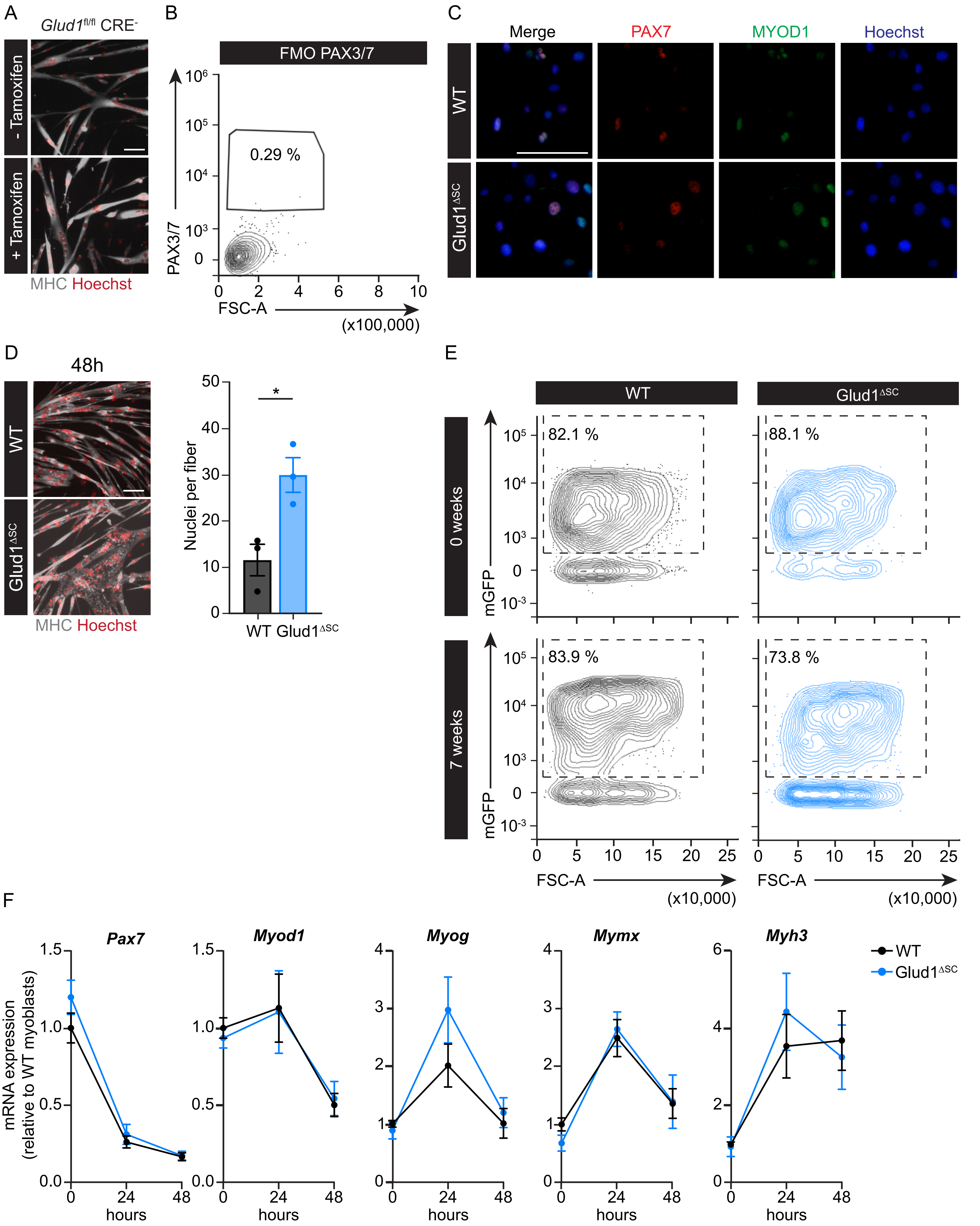

A
Glud1fl/fl CRE-
- Tamoxifen
+ Tamoxifen
MHC Hoechst
B
FMO PAX3/7
0.29 %
PAX3/7
FSC-A
(x100,000)
C
Merge
PAX7
MYOD1
Hoechst
WT
Glud1ΔSC
D
48h
WT
Glud1ΔSC
MHC Hoechst
Nuclei per fiber
*
WT Glud1ΔSC
E
WT
Glud1ΔSC
0 weeks
7 weeks
82.1 %
88.1 %
83.9 %
73.8 %
mGFP
FSC-A
(x10,000)
F
Pax7
Myod1
Myog
Mymx
Myh3
mRNA expression (relative to WT myoblasts)
hours
WT
Glud1ΔSC

**Figure S3. Loss of *Glud1* impairs myoblast self-renewal capacity and promotes fusion without affecting the core transcriptional regulation of MuSC differentiation. Related to Figure 2, Figure 3, and Figure 4.**

(A) Representative images of myosin heavy chain (MHC) immunofluorescent staining (grey, MHC; red, Hoechst; scale bar 100 µm) in myotubes derived from *Pax7-CreER*$^{T2-/-}$-*Glud1*$^{fl/fl}$ (*Glud1*$^{fl/fl}$CRE$^{-}$) myoblasts with (+Tamoxifen) or without tamoxifen (-Tamoxifen) treatment to induce recombination at the myoblast stage. (B) Representative flow cytometric analysis showing fluorescence minus one (FMO) control of Pax3/7$^{+}$ staining in myoblasts. Related to Figure 2F. (C) Representative images of PAX7 and MYOD1 immunofluorescent staining (red, PAX7; green, MYOD1; blue, Hoechst; scale bar 100 µm) in WT and *Glud1*$^{\Delta SC}$ myoblasts. Related to Figure 2G. (D) Representative images (left) of myosin heavy chain (MHC) immunofluorescent staining (grey, MHC; red, Hoechst; scale bar 100 µm) in WT and *Glud1*$^{\Delta SC}$ myotubes after acute tamoxifen treatment and quantification (right) of the nuclei per myotube 48 h after the induction of differentiation. (E) Representative flow cytometric analysis of mGFP$^{+}$ cells as a proportion of PAX7$^{+}$ cells in *Glud1*$^{WT-mG}$ and *Glud1*Δ$^{SC-mG}$ mice 1 week after tamoxifen treatment (0 weeks) and after wheel running (7 weeks). Related to Figure 3C. (F) Gene expression analysis by qPCR of the canonical myogenic genes *Pax7, Myod1, Myog, Mymx,* and *Myh3* in WT and *Glud1*$^{\Delta SC}$ cells at 3 different steps during differentiation before (0 h), during (24 h), and upon completion of (48 h) differentiation. n= 5 independent experiments. Related to Figure 4. Graph bars represent mean $\pm$ SEM. Dots represent independent experiments. Student's T- test (two tailed unpaired) in (D). Two-way ANOVA with Sidak's multiple comparison test in (F). None of the comparisons were significant in (F). (*p < 0.05, **p < 0.01, ***p < 0.001).

**Figure S4**

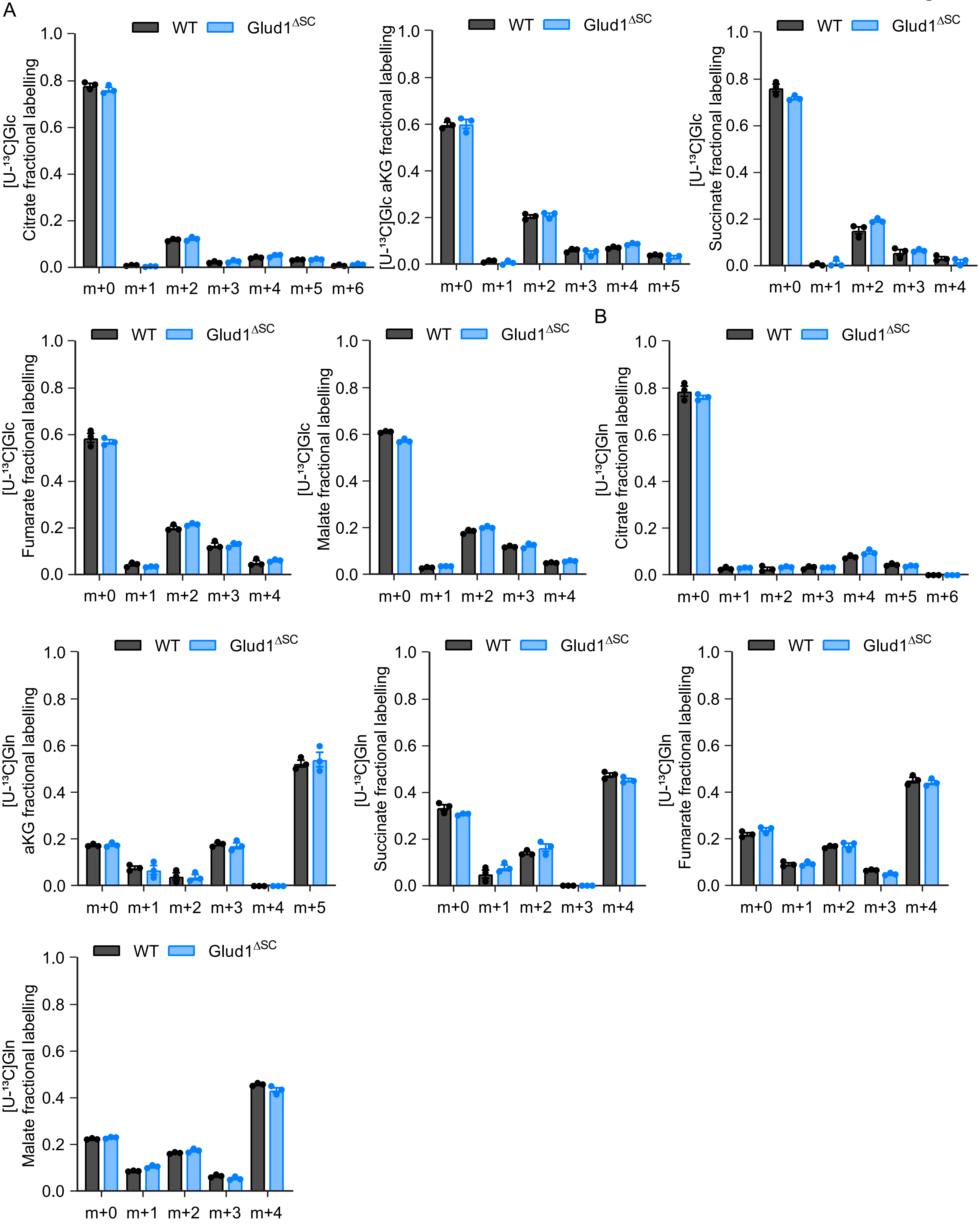

A
WT
Glud1ΔSC
[U-13C]Glc Citrate fractional labelling
1.0
0.8
0.6
0.4
0.2
0.0
m+0
m+1
m+2
m+3
m+4
m+5
m+6
[U-13C]Glc aKG fractional labelling
[U-13C]Glc Succinate fractional labelling
[U-13C]Glc Fumarate fractional labelling
[U-13C]Glc Malate fractional labelling
B
[U-13C]Gln Citrate fractional labelling
[U-13C]Gln aKG fractional labelling
[U-13C]Gln Succinate fractional labelling
[U-13C]Gln Fumarate fractional labelling
[U-13C]Gln Malate fractional labelling

**Figure S4. Mass distribution vectors in TCA cycle metabolites during [U-$^{13}$C]glucose and [U-$^{13}$C]glutamine tracing. Related to Figure 5.**

(A) Mass distribution vector (MDV) of citrate, alpha-ketoglutarate (aKG), succinate, fumarate and malate during [U-$^{13}$C]Glc tracing in WT and *Glud1*$^{\Delta SC}$ myoblasts. (B) MDV of citrate, aKG, succinate, fumarate, and malate during [U-$^{13}$C]Gln tracing in WT and *Glud1*$^{\Delta SC}$ myoblasts. Bar graphs represent mean ± SEM. Each dot represents a biological replicate.

**Figure S5**

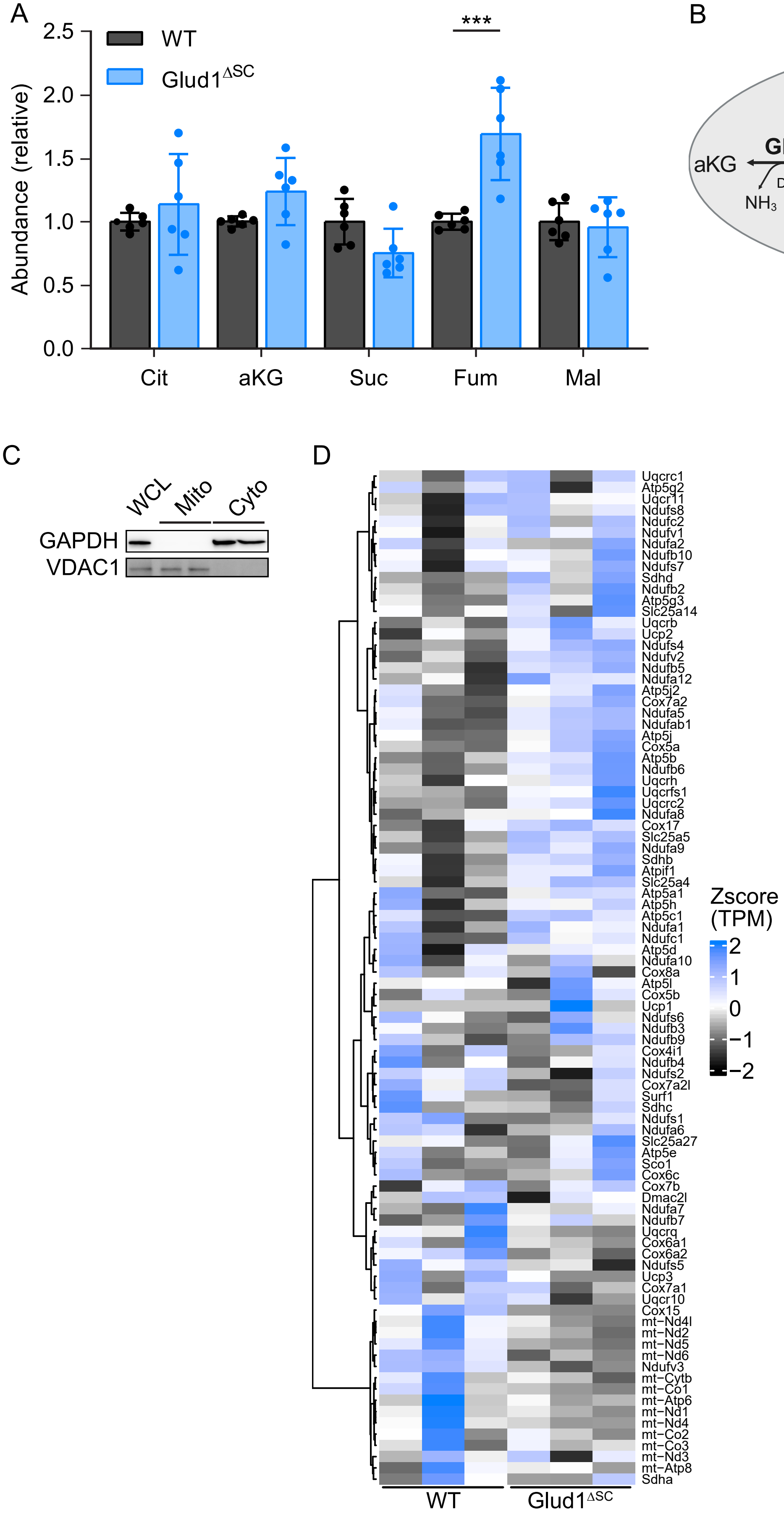
A
2.5
2.0
1.5
1.0
0.5
0.0
Abundance (relative)
WT
Glud1^ΔSC
***
Cit
aKG
Suc
Fum
Mal
B
Glutamine
Cytoplasm
Mitochondria
Glutamine
GLS
Deamidation
NH3
GLUD1
aKG
Glutamate
Deamination
NH3
GOT2,
GPT2, BCAT2
aKG
Transamination
alpha-
ketoacid
amino
acid
GOT1,
GPT1, BCAT1
Glutamate
aKG
Transamination
alpha-
ketoacid
amino
acid
C
WCL
Mito
Cyto
GAPDH
VDAC1
D
Uqcrc1
Atp5g2
Uqcr11
Ndufs8
Ndufc2
Ndufv1
Ndufa2
Ndufb10
Ndufs7
Sdhd
Ndufb2
Atp5g3
Slc25a14
Uqcrb
Ucp2
Ndufs4
Ndufv2
Ndufb5
Ndufa12
Atp5j2
Cox7a2
Ndufa5
Ndufab1
Atp5j
Cox5a
Atp5b
Ndufb6
Uqcrh
Uqcrfs1
Uqcrc2
Ndufa8
Cox17
Slc25a5
Ndufa9
Sdhb
Atpif1
Slc25a4
Atp5a1
Atp5h
Atp5c1
Ndufa1
Ndufc1
Atp5d
Ndufa10
Cox8a
Atp5l
Cox5b
Ucp1
Ndufs6
Ndufb3
Ndufb9
Cox4i1
Ndufb4
Ndufs2
Cox7a2l
Surf1
Sdhc
Ndufs1
Ndufa6
Slc25a27
Atp5e
Sco1
Cox6c
Cox7b
Dmac2l
Ndufa7
Ndufb7
Uqcrq
Cox6a1
Cox6a2
Ndufs5
Ucp3
Cox7a1
Uqcr10
Cox15
mt−Nd4l
mt−Nd2
mt−Nd5
mt−Nd6
Ndufv3
mt−Cytb
mt−Co1
mt−Atp6
mt−Nd1
mt−Nd4
mt−Co2
mt−Co3
mt−Nd3
mt−Atp8
Sdha
WT
Glud1^ΔSC
Zscore
(TPM)
2
1
0
−1
−2

**Figure S5. Evaluation of mitochondrial pathways upon the loss of *Glud1.* Related to Figure 5.**

(A) Relative abundance of the TCA cycle metabolites citrate (Cit), alpha-ketoglutarate (aKG), succinate (Suc), fumarate (Fum), and malate (Mal) in WT and *Glud1*$^{\Delta SC}$ myoblasts. (B) Schematic representation of the cytosolic and mitochondrial glutamate metabolism pathways showing the mitochondrial (GOT2, GPT2, BCAT2) and cytosolic (GOT1, GPT1, BCAT1) aminotransferases and the mitochondrial glutamate deaminase (GLUD1), which does not have a cytosolic counterpart. (C) Western blot analysis of the cytosolic protein GAPDH and the mitochondrial protein VDAC1 in the cytosolic fraction (Cyto), mitochondrial fraction (Mito), and whole cell lysates (WCL) of myoblasts. (D) Expression of electron transport chain genes (WikiPathways) in WT and *Glud1*$^{\Delta SC}$ myoblasts. Expression is evaluated using transcript per million (TPM) z-scored for each gene. Bar graphs represent the mean $\pm$ SEM. Each dot represents a biological replicate. Two-way ANOVA with Sidak's multiple comparison test in (A). (*$p < 0.05$, **$p < 0.01$, ***$p < 0.001$, ****$p<.0001$).

**Figure S6**

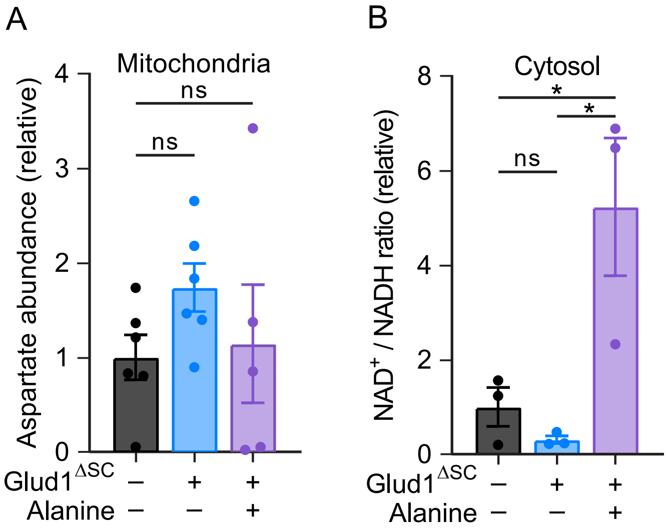
A
Mitochondria
Aspartate abundance (relative)
4
3
2
1
0
ns
ns
Glud1$^{\Delta SC}$ − + +
Alanine − − +
B
Cytosol
NAD$^+$ / NADH ratio (relative)
8
6
4
2
0
*
*
ns
Glud1$^{\Delta SC}$ − + +
Alanine − − +

**Figure S6. Alanine supplementation rescues MAS deficiency and imbalanced fusion in *Glud1* deficient MuSCs. Related to Figure 6.**

(A) Relative mitochondrial aspartate levels in WT and *Glud1*$^{\Delta SC}$ myoblasts grown in growth media with or without 0.2 mM alanine supplementation. (B) Relative cytosolic $NAD^+$ / NADH ratio in WT and *Glud1*$^{\Delta SC}$ myoblasts grown in growth media with or without 0.2 mM alanine supplementation. Bar graphs represent the mean $\pm$ SEM. Each dot represents a biological replicate. One-way ANOVA with Dunnett's multiple comparison test in (A). One-way ANOVA with Tukey's multiple comparison test in (B). (*$p < 0.05$, **$p < 0.01$, ***$p < 0.001$, ****$p < .0001$).

**Table S1.**

| Classic media | | |
|---|---|---|
| Compound | Proliferation media (mM) | Differentiation media (mM) |
| Glucose | 4.11 | 24.50 |
| Glycine | 0.16 | 0.39 |
| L-Alanine | 0.04 | 0.00 |
| L-Arginine hydrochloride | 0.52 | 0.39 |
| L-Asparagine-H2O | 0.04 | 0.00 |
| L-Aspartic acid | 0.04 | 0.00 |
| L-Cysteine | 0.14 | 0.20 |
| L-Glutamic Acid | 0.04 | 0.00 |
| L-Glutamine | 1.59 | 3.92 |
| L-Histidine hydrochloride-H2O | 0.10 | 0.20 |
| L-Isoleucine | 0.25 | 0.79 |
| L-Leucine | 0.28 | 0.79 |
| L-Lysine hydrochloride | 0.30 | 0.78 |
| L-Methionine | 0.07 | 0.20 |
| L-Phenylalanine | 0.13 | 0.39 |
| L-Proline | 0.04 | 0.00 |
| L-Serine | 0.16 | 0.39 |
| L-Threonine | 0.25 | 0.78 |
| L-Tryptophan | 0.02 | 0.08 |
| L-Tyrosine disodium salt dihydrate | 0.12 | 0.39 |
| L-Valine | 0.25 | 0.79 |

**Table S1. Comparison of the glucose and amino acid concentrations present in the classic proliferation and differentiation media for MuSCs. Related to Figure 1 and Figure S1**

**Table S2.**

| Gene | Forward | Reverse |
|---|---|---|
| *Pax7* | CAGTGTGCCATCTACCCATGCTTA | GGTGCTTGGTTCAAATTGAGCC |
| *Myod1* | TGGGATATGGAGCTTCTATCGC | GGTGAGTCGAAACACGGATCAT |
| *MyoG* | CATCCAGTACATTGAGCGCCTA | GAGCAAATGATCTCCTGGGTTG |
| *Mymx* | GTTAGAACTGGTGAGGAGGAG | CCATCGGGAGCAATGGAA |
| *Myh3* | ACATCTCTATGCCACCTTCGCTAC | GGGTCTTGGTTTCGTTGGGTAT |

**Table S2. Sequences of primers used for RT-PCR. Related to Figure 4 and Figure S6**
